# Analysis of Polycerate Mutants Reveals the Evolutionary Co-option of *HOXD1* to Determine the Number and Topology of Horns in Bovidae

**DOI:** 10.1101/2020.11.04.363069

**Authors:** Aurélie Allais-Bonnet, Aurélie Hintermann, Marie-Christine Deloche, Raphaël Cornette, Philippe Bardou, Marina Naval-Sanchez, Alain Pinton, Ashleigh Haruda, Cécile Grohs, Jozsef Zakany, Daniele Bigi, Ivica Medugorac, Olivier Putelat, Ockert Greyvenstein, Tracy Hadfield, Slim Ben Jemaa, Gjoko Bunevski, Fiona Menzi, Nathalie Hirter, Julia M. Paris, John Hedges, Isabelle Palhiere, Rachel Rupp, Johannes A. Lenstra, Louisa Gidney, Joséphine Lesur, Renate Schafberg, Michael Stache, Marie-Dominique Wandhammer, Rose-Marie Arbogast, Claude Guintard, Amandine Blin, Abdelhak Boukadiri, Julie Riviere, Diane Esquerré, Cécile Donnadieu, Coralie Danchin-Burge, Coralie M Reich, David Riley, Este van Marle-Koster, Noelle Cockett, Benjamin J. Hayes, Cord Drögemüller, James Kijas, Eric Pailhoux, Gwenola Tosser-Klopp, Denis Duboule, Aurélien Capitan

**Author notes:** These authors contributed equally to this work (Aurélie Allais-Bonnet, Aurélie Hintermann). Corresponding authors: Aurélien Capitan, Université Paris-Saclay, INRAE, AgroParisTech, GABI, 78350 Jouy-en-Josas, France, Denis Duboule, Department of Genetics and Evolution, University of Geneva, 1211, Geneva 4, Switzerland, Swiss Cancer Research Institute, EPFL, Lausanne, Suisse, College de France, Paris, France.

## Abstract

In the course of evolution, pecorans (i.e. higher ruminants) developed a remarkable diversity of osseous cranial appendages, collectively referred to as ‘headgear’, which likely share the same origin and genetic basis. However, the nature and function of the genetic determinants underlying their number and position remain elusive. Jacob and other rare populations of sheep and goats, are characterized by polyceraty, the presence of more than two horns. Here, we characterize distinct *POLYCERATE* alleles in each species, both associated with defective *HOXD1* function. We show that haploinsufficiency at this locus results in the splitting of horn bud primordia, likely following the abnormal extension of an initial morphogenetic field. These results highlight the key role played by this gene in headgear patterning and illustrate the evolutionary co-option of a gene involved in the early development of bilateria to properly fix the position and number of these distinctive organs of Bovidae.

## Introduction

In pecorans, successive environmental and behavioural adaptations favoured the emergence and sometimes the secondary loss of a variety of headgear, as exemplified by bovid horns, cervid antlers, giraffid ossicones or antilocaprid pronghorns (Davis et al. 2011; Wang et al. 2019). As different as they are, these iconic organs share both a common cellular origin and a minimal structural organisation: they derive from neural crest stem cells and consist of paired structures, located on the frontal bones and composed of a bony core covered by integument (Davis et al. 2011; Wang et al. 2019) (**Fig. 1, Suppl. Fig. 1**). While the development and evolution of headgear is a long-standing question, the underlying molecular and cellular mechanisms have been difficult to study, mostly because the patterning and differentiation of headgear progenitor cells occur early during embryogenesis (Lincoln 1973; Allais-Bonnet et al. 2013) and involve hundreds of genes (Wang et al. 2019).

**Figure 1.**
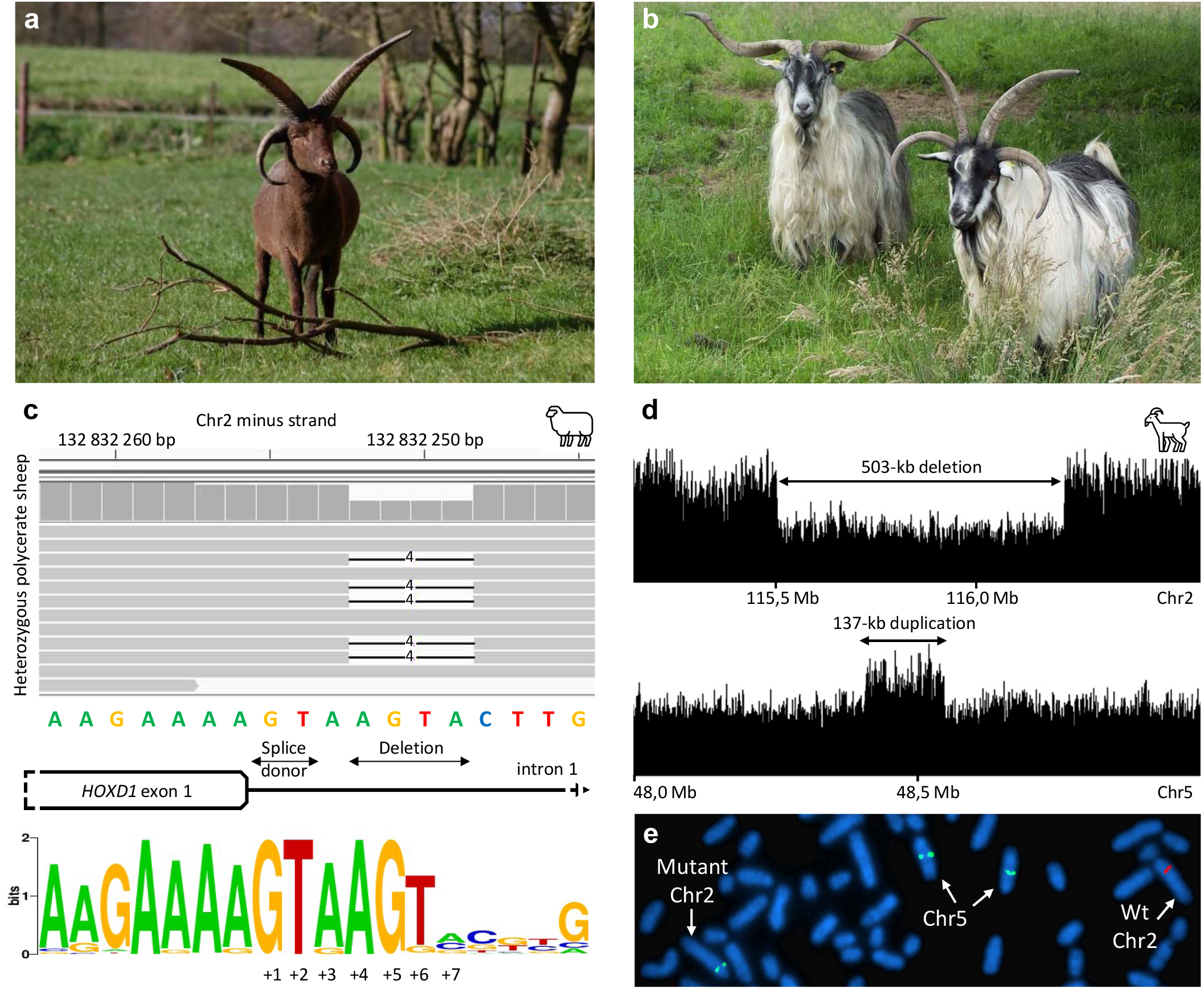
Polyceraty in sheep and goats and candidate genetic variants. a) Polycerate Manx Loaghtan ram. b) Wildtype and polycerate male goats from a local German population. These individuals represent the most common phenotype. Polycerate animals with asymetric horns and partial fusion of lateral horns are also regularly observed. c) A 4bp deletion causing polyceraty in sheep. Integrative Genome Viewer (IGV) screenshot with the localization of the variant with respect to HOXD1. Below is a graphical representation of nucleotide conservation at the exon 1-intron junction across 103 sarcopterigian and tetrapod species. d) Plot of read coverage in a heterozygous polycerate goat animal carrying a deletion of 503 kb downstream the *HOXD* gene cluster on Chr2 and a duplication of 137 kb on Chr5. e) FISH-mapping in a heterozygous polycerate goat with BAC clones corresponding to the region deleted in Chr2 (labeled in red) and to the segment of Chr5 inserted at the deletion site (labeled in green). Magnification: X1000. Sheep and goat icons were made by ‘Monkik’ from www.thenounproject.com.

In this context, natural mutations affecting headgear number, shape or position, such as the polycerate (multi-horned) phenotype occurring in small ruminants (**Fig. 1a, b,** OMIA 000806-9940), offer a valuable alternative (Capitan et al. 2012). Polyceraty was already observed in the oldest ovine remains from Çatalhöyük, Turkey (ca 6000 BCE (Epstein 1971; Putelat 2005)) and this dominant phenotype currently segregates in several sheep breeds around the world. Even though the corresponding locus was mapped in seven distinct populations to the same region of chromosome 2 (Chr2), it has not yet been identified (Greyvenstein et al. 2016; He et al. 2016; Kijas et al. 2016; Ren et al. 2016). In contrast, polycerate goats are found only sporadically in the Alps and have not been subject to any genetic studies thus far. The oldest record dates back from 1786, when the French Queen Marie-Antoinette imported a four-horned billy-goat from the city of Bulle, Switzerland to her model farm (Heitzmann 2006).

We set up to determine the genetic bases of these conditions in sheep and goats and, in this study, we show that polyceraty in Bovidae is due either to a four-base-pair deletion affecting the splicing of the *Hoxd1* gene in sheep, or to the deletion of a large regulatory region controling the same gene, in goats. These results thus illustrate the evolutionary co-option of this gene normally involved in early development to help determine the position and number of horns. They also show that comparable phenotypes observed in distinct species and selected and maintained for a long time are caused by the mis-regulation of the same gene.

## Results and Discussion

### Characterisation of *POLYCERATE* Mutations in Sheep and Goats

To identify the molecular determinants of polyceraty, we re-analysed the Illumina OvineHD Beadchip genotyping data (600 k SNPs) of 111 case and 87 control sheeps generated by two previous studies (Greyvenstein et al. 2016; Kijas et al. 2016) (**Suppl. Tables 1 and 2**). Assuming autosomal dominant inheritance and genetic homogeneity in the three breeds investigated, we fine-mapped the ovine *POLYCERATE* locus between positions 132,717,593 and 133,151,166 -bp on Chr2 (**Suppl. Fig. 2**). By comparing whole genome sequences of 11 polycerate specimens and 1’179 controls representing the world-wide sheep diversity, we identified a single candidate variant in this interval: a four-nucleotide deletion located at position +4 to +7 bp after exon 1 of the *HOXD1* gene (g.132,832,249_132,832,252del; **Fig. 1c**). Genotyping of this variant in 236 animals from eight populations containing polycerate specimens showed a perfect genotype to phenotype association (**Suppl. Tables 3 and 4**). Moreover, cross-species alignments revealed that the +4 and +5 nucleotides are conserved amongst 103 sarcopterigian and tetrapod species, supporting an important role in the splicing of *HOXD1* precursor RNAs (**Fig. 1c** and **Suppl. Table 5**).

We next mapped the caprine *POLYCERATE* locus to a 542 kb large region orthologous to that of the ovine locus (Chr2:115,143,037-115,685,115 bp on ARS1 assembly (Bickhart et al. 2017); **Suppl. Fig. 2**), by using a panel of 35 polycerate and 51 two-horned goats obtained from eight European populations and genotyped with the Illumina GoatSNP50 BeadChip (Tosser-Klopp et al. 2014) (**Suppl. Table 6**). Within this interval, we identified 36 private heterozygous variants in one heterozygous polycerate goat *versus* 1’160 control individuals (**Suppl. Table 7**). Genotyping of five case-control pairs from distinct breeds reduced the list of candidates to 15 short variants, affecting genomic regions not conserved amongst 103 eutherian mammals, as well as a rare type of structural variation located 57 kb downstream of the *HOXD1* 3’UTR (**Suppl. Tables 7 and 8**). The latter involved the translocation of 137 kb from Chr5 to Chr2 by means of a circular intermediate (Durkin et al. 2012) and the deletion of 503 kb from the insertion site (g.115,652,290_116,155,699delins137kb; **Fig. 1d, e** and **Suppl. Fig. 3**). Consequently, the mutant chromosome lacked the *MTX2* gene and carried an exogenic copy of both *RASSF3* and the first ten exons of *GNS*. Genotyping of this variant in 77 case and 355 control goats originating from 24 distinct populations revealed a 100 percent association between polyceraty and heterozygosity for the large insertion/deletion (**Suppl. Table 9** and **Suppl. Fig. 4**). Homozygous mutants were not detected in our panel, whereas at least 14 polycerate animals were born from polycerate pairs of parents (binomial p = 3.4 x 10^-3^; **Suppl. Note 1**). Because the knockdown of *Mtx2* in zebrafish is embryonic lethal at gastrulation (Wilkins et al. 2008) and newborn mice homozygous for a deletion including *Mtx2* were never scored (binomial p = 5.7 x 10^-6^; **Suppl. Note 1**), we concluded that homozygosity at the goat *POLYCERATE* locus is an early lethal condition.

### Remote *Hoxd1* regulation in Transgenic Mice

These mapping studies identified the *HOXD* gene cluster as being involved in the polycerate phenotype in both sheep and goats. This cluster contains nine homeobox genes encoding transcription factors involved in the organisation of the body plan during embryogenesis (Krumlauf 1994). Both their timing of activation and their domains of expression are determined by their respective positions along the gene cluster (Kmita and Duboule 2003). Accordingly, the mouse *Hoxd1* gene is expressed very early on and in the most rostral part of the embryo (**Fig. 2a**). In rodents, *Hoxd1* is expressed in crest cell-derived head structures (Frohman and Martin 1992), which made this gene a particularly interesting candidate for polyceraty. Also, a DNA sequence conserved only amongst pecorans species carrying headgear was identified 15 kb downstream of *HOXD1* (“HCE” in (Wang et al. 2019)). This sequence, however, is not included in the large insertion-deletion observed in polycerate goats.

**Figure 2.**
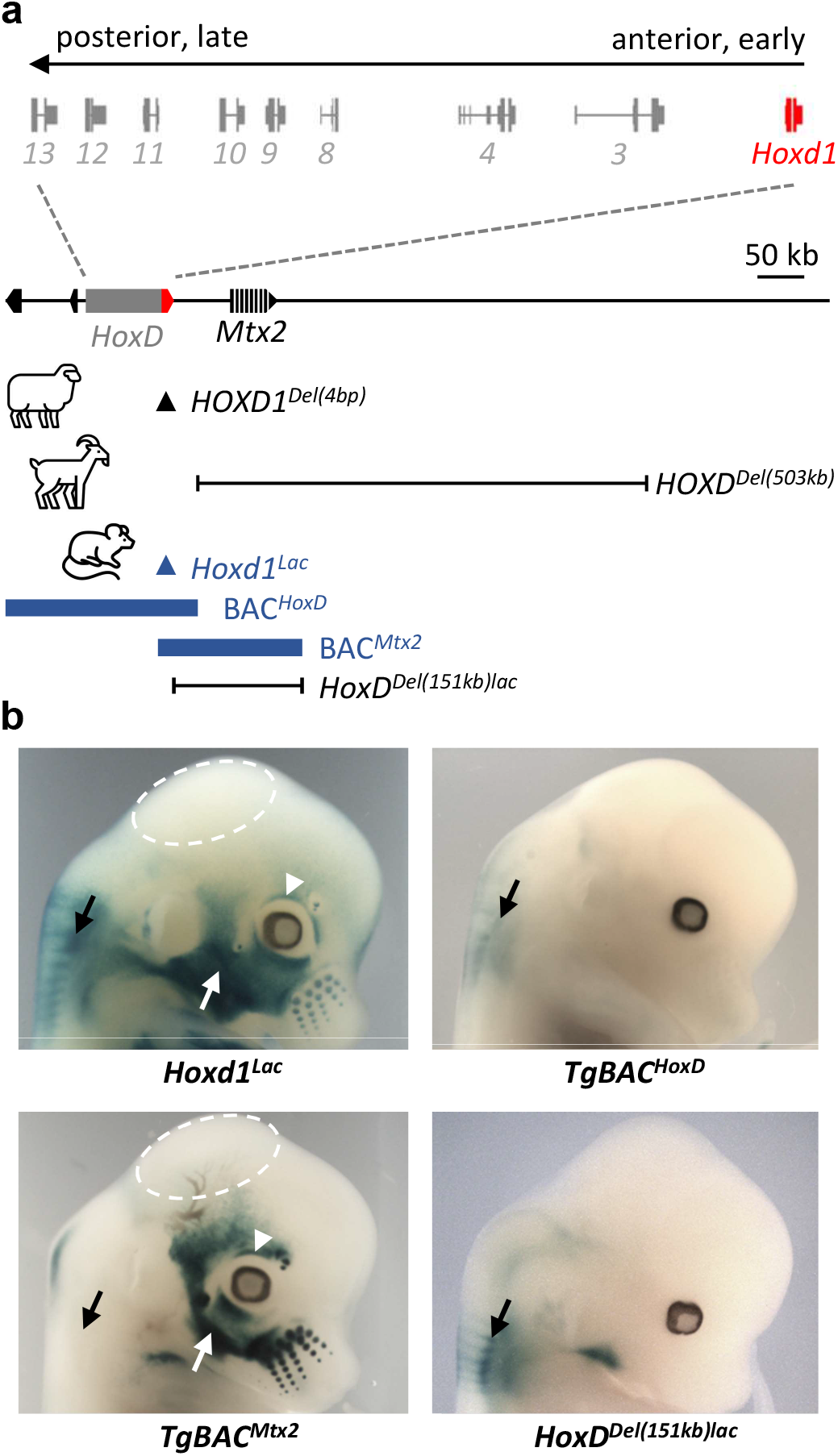
Regulation of *Hoxd1* expression pattern in crest cell-derived head structures in mouse. a) On top is the structure of the mouse *HoxD* gene cluster with arrows showing the timing and localisation of gene expression along the body axis during development. The position of *Hoxd1* is highlighted in red. Below is a 1 Mb view of the locus, with *Hoxd1* in red as well as the relative position of the *POLYCERATE* variants in sheep (black arrowhead) and goat (black line). Below are depicted the various murine alleles, with the lacZ insertion in *Hoxd1* (blue arrowhead), the two BAC clones (thick blue lines) and the engineered deletion (black line). b) Fetal heads of E12.5-E13.5 mouse fetuses after X-gal staining. The dashed circle highlights the absence of *Hoxd1* expression in the crown (corresponding to the localization of hornbuds in Bovidae), whereas the surrounding dermal cells are positive. The conservation of *Hoxd1* expression in the back of the neck (black arrows) contrasts with the presence/absence of expression in the facial muscle precursors (white arrows) and in the eyelids (arrowhead). The comparison between the four strains indicate that *Hoxd1* expression in all these cranial derivatives is controlled by regulatory elements located in a region orthologous to the proximal portion of the segment deleted in polycerate goats.

We assessed whether the deletion present in goat may impact the expression of *HOXD1* in cranial crest cells by looking at a series of modified mouse strains either carrying transgenes or where a targeted deletion was induced at the orthologous locus (see Methods). First, the wide presence of cells expressing *Hoxd1* both in the face and in the cranial derma, the latter being of crest cell origin, was detected in fetuses with a targeted integration of lacZ sequences into the *Hoxd1* gene (**Fig. 2b**, *Hoxd1^Lac^*). Expression was however not scored in the crown region (**Fig. 2b**, dashed circle), the area corresponding to that of horn bud differentiation in Bovidae (Dove 1935; Capitan et al. 2011). Instead, *Hoxd1* was expressed abundantly in other regions of the head including the eyelids (**Fig. 2b**, white arrow and arrowhead), an observation consistent with the abnormal upper eyelids and eyebrows often detected in ovine and caprine polycerate animals (**Suppl. Fig. 5-7)**.

We next tried to localize the underlying regulatory elements by using transgenic BACs with lacZ sequences introduced within *Hoxd1*. A BAC covering the *HoxD* cluster itself did not show any expression in the head, suggesting that regulatory sequences are not located in the gene cluster (**Fig. 2b**, *TgBAC^HoxD^*). In contrast, a transgenic BAC extending in the region upstream of *Hoxd1* and including *Mtx2* gave a staining similar to *Hoxd1^Lac^* (**Fig. 2b**, *TgBAC^Mtx2^*), indicating that regulatory sequences were located upstream *Hoxd1*, in a region including and surrounding *Mtx2*. The latter result was controlled by using an engineered 151 Kb deletion of a largely overlapping region, including a lacZ reporter gene, which expectedly abrogated *Hoxd1* expression in cranial cellular populations (**Fig. 2b,** *HoxD^Del(151kb)lac^*). As a positive control for the lacZ reporter system, expression of *Hoxd1* in neural derivatives driven by sequences within the *HoxD* cluster was scored, as expected (**Fig. 2b**, black arrows). These analyses in mice demonstrated that regulatory sequences driving *Hoxd1* expression in the head are located in a region largely comprised within the deletion determined in goats as causative of polyceraty, further suggesting that the latter deletion abrogates *HOXD1* expression in goat fetuses.

### Expression of *HOXD1* in Pecoran Fetuses

To investigate whether the abence of mouse *Hoxd1* expression in the crown region of the head was also observed in Pecoran embryos, we isolated heterozygous polycerate and wild type fetuses both at 70 dpc (days *post-coïtum*) in goat and at 76 dpc in sheep, two stages where eyelids are fully grown and horn buds can be distinguished (**Suppl. Fig. 8**). After micro-dissection and Reverse Transcription quantitative PCR (RT-qPCR), we noticed that in wild type fetuses of both species, *HOXD1* expression was significantly lower in horn buds than in surrounding tissues (**Fig. 3a, b**), reminiscent of the weak -if any-expression of *Hoxd1* observed in a comparable region in the mouse. In heterozygous mutant goat fetuses, however, *HOXD1* RNA levels were equally low in all three samples (**Fig. 3a**), re-enforcing the idea that the caprine *POLYCERATE* variant negatively affects the expression of *HOXD1*.

**Figure 3.**
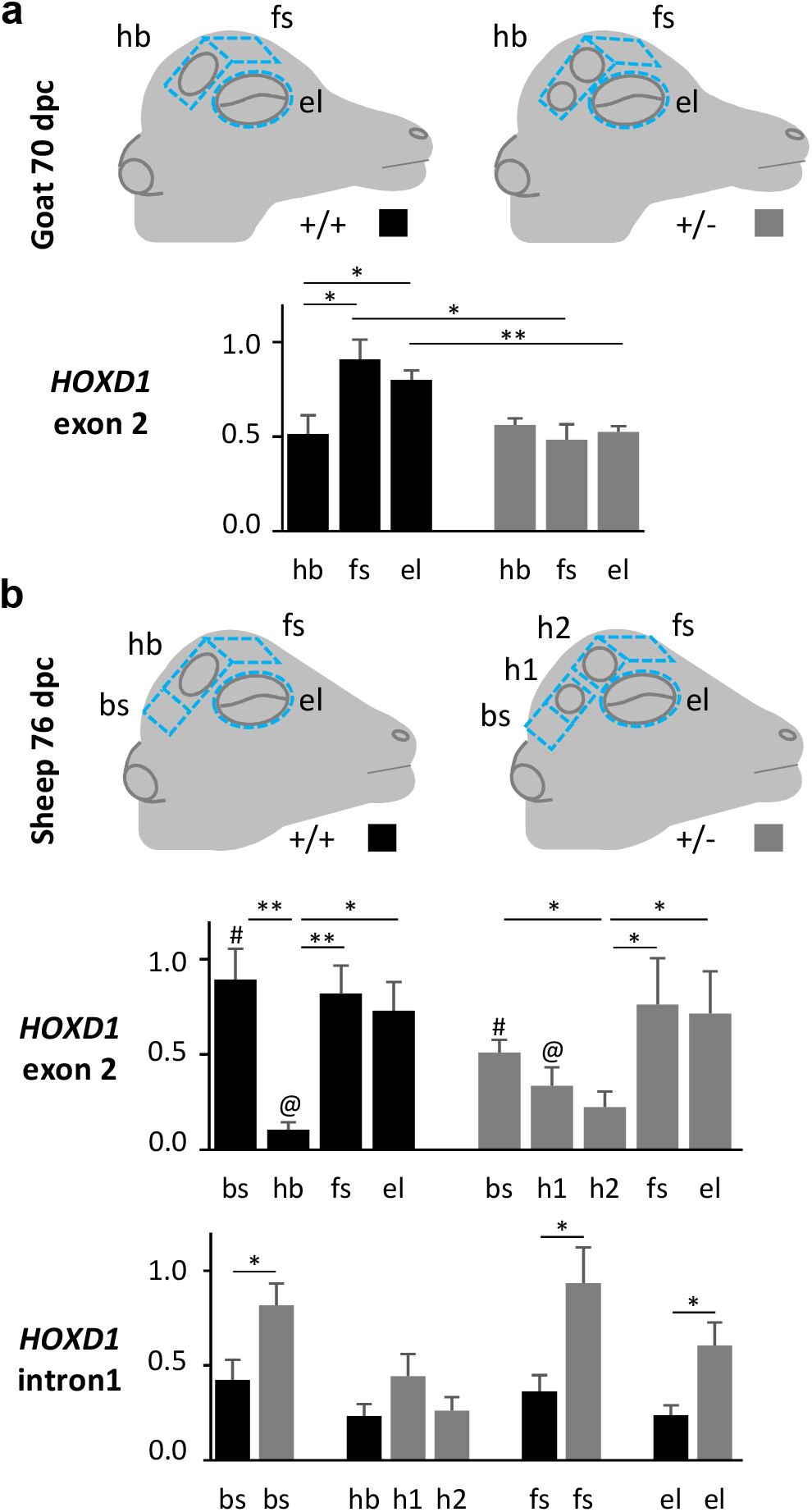
RT-qPCR gene expression analyses in sheep and goat foetuses. a, b) Schemes of the tissues sampled at stage 70 dpc in goat (a) and 76 dpc in sheep (b) in four control (+/+) and four heterozygous (+/-) polycerate fetuses within each species. bs: skin from the back of the head; hb: skin from the hornbud; h1: skin from the lower horn bud; and h2: skin from the upper horn bud in polycerate specimens; fs: frontal skin; el: eyelids. RT-qPCR gene expression analyses in these tissues are shown below (means and standard errors of the means). *: p<0.05, **: p<0.01 (Welch two sample t-test with the alternative hypothesis that the means are not equal). For the sake of clarity, the symbols # and @ were also used to show significant differences (p <0.05) between distant bars.

In sheep, despite some variation due to slight differences in sampling (**Fig. 3b,** upper histogram and methods), heterozygous mutants for the four-base-pair deletion adjacent to the splice donor site displayed RNA amounts roughly similar to control specimens when primers targeting the second exon of the gene were used. However, RT-qPCR with intronic primers revealed significant intron retention in all mutant tissues but horn buds, where expression was likely too low (**Fig. 3b** lower histogram). Intron retention is predicted to result in a non-functional protein, truncated two residues after the last amino acid encoded by exon 1 and thus lacking the DNA binding peptide (**Suppl. Fig. 9**). Therefore, both *POLYCERATE* variants appear to reduce the amount of functional *HOXD1* RNAs in the horn bud region, likely leading to a loss of boundary condition and an extension of the cellular field permissive for horn bud development. This extension would elongate the bud region sufficiently to split it into two separate organs.

### Morphometric Analyses and Topology of the Horn Field

To substantiate this hypothesis, we analysed variations in horn topology in 61 ovine and 19 caprine skulls from various populations using 3D geometric morphometrics (**Suppl. Table 10**). We performed Principal Component Analysis (PCA) using 116 landmarks and sliding semi-landmarks after removing non-shape variation (see Methods). We then plotted the first principal components (PCs) to visualize the specimen distribution in the morphospace (**Fig. 4a, Suppl. Fig. 10-12** and **Suppl. Table 11**). The first two axes represented 35.8% and 23.3% of the total variance and distinguished the phenotype and species categories, respectively. Along the first axis, we individualized three sub groups of polycerate specimens in sheep, based on the distances between lateral horns (dlh, **Fig. 4a**). Of note, the group displaying the largest dlh (i.e. that with the most negative values along the x axis) had no equivalent in goat, possibly due to early lethal homozygosity (see above).

**Figure 4.**
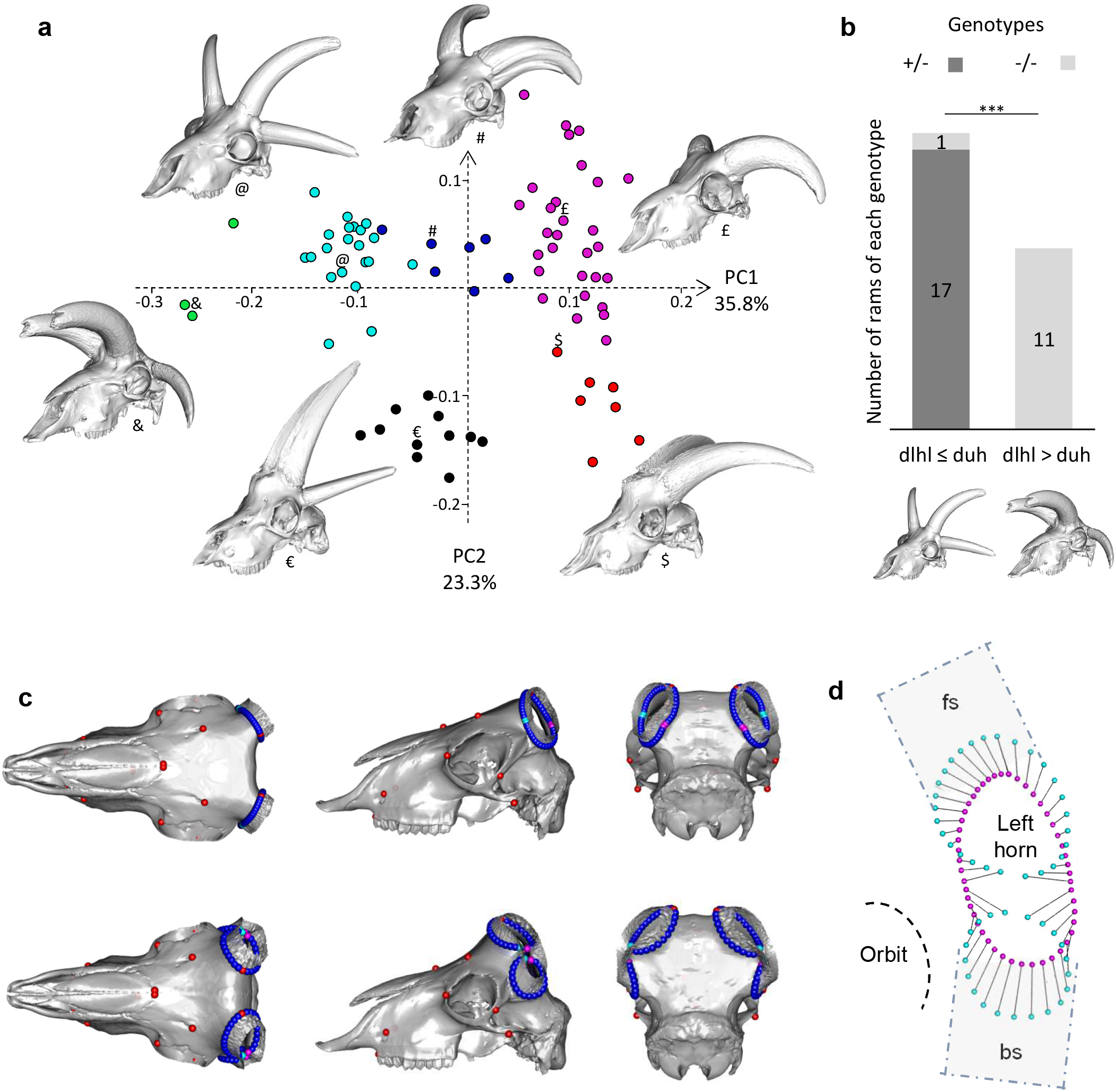
Results of three-dimensional geometric morphometric analyses of 61 ovine and 19 caprine skulls. a) Distribution of the specimens along the first two axes of the PCA. The proportion of variance explained by the main principal components is indicated on each axis. Green dots: polycerate sheep with a distance between lateral horns (dlh) larger than the distance between upper horns (duh); light blue: polycerate sheep with a dlh≤duh; blue: polycerate sheep with at least two lateral horns partially fused at their basis; purple: wild type sheep; black: polycerate goats; and red: wild type goats. Representative specimens illustrate each cluster and symbols are used to indicate their respective locations in the PCA analysis (see **Suppl. Fig. 10** for intraspecies analyses and further information). b) Number of heterozygous (+/-) and homozygous (-/-) polycerate rams amongst groups of live animals with different dlhl (dlh on the left side) and duh relative sizes (see **Suppl. Table 12** for further information); ***: p-value= 3.5 x 10^-7^ (Fisher’s exact test). c) Theoretical shapes associated with the maximum (upper three) and minimum values (lower three) of PC1 axis for a sheep skull. Red dots correspond to anatomical landmarks while the other dots correspond to sliding semi-landmarks; light blue and purple dots highlight the sites of division of lateral horns. d) Shape differences for the sliding semi-landmarks located at the basis of the left horn. Light blue and purple dots correspond to the maximum and minimum values of PC1 axis, respectively. Dashed squares indicate the estimated position of dissected tissues in **Fig. 3** (bs: skin from the back of the head; fs: frontal skin) in which *HOXD1* expression was observed in fetuses.

We looked at the association between genotypes and horn implantation within polycerate animals by measuring the distances both between the lateral horns on the left side of the skull (dlhl) and between the upper horns (duh) in 29 rams (**Suppl. Table 12**). We found a significant difference in the proportions of homozygous and heterozygous specimens in animals with dlhl≤duh *versus* dlhl>duh (**Fig. 4b**) and no heterozygous animal was found to have dlhl>duh. We computed the theoretical skull shape at the maximum and minimum of PC1 axis (**Fig. 4c**) and the corresponding vectors of deformation (**Fig. 4d**). The results obtained were consistent with a splitting of horn buds in polycerate animals. This splitting always occurred along the major axis of the ellipse formed by the wild type horn bud, with an extension of the hemi-horn buds in an area where *HOXD1* expression was detected in wild type specimens (**Fig. 4d** and above). In homozygous animals, the new cellular field was likely larger than in heterozygous, leading to a clearer separation of hemi-horns, whereas heterozygous specimens often displayed partially fused organs. This is markedly different from the production of additional horns, as observed in subspecies of *Tetracerus quadricornis* (Groves 2003) (**Suppl. Fig. 13**).

## Conclusions

From these results, we conclude that pecorans have an intrinsic capacity to induce hornbuds within a presumptive head territory. This capacity appears to be associated with the non-expression of the *HOXD1* transcription factor, which is present in surrounding cells and may delimit this field, a function somewhat distinct from the ancestral role of *Hox* genes during development (Krumlauf 1994). Various haploinsufficient conditions lead to the extension of this territory, a condition fully achieved in complete absence of *HOXD1*. While a weak extension of this morphogenetic field leads to the growth of twin horns, fused at their bases, a full extension induces the complete splitting of the horn bud, thus generating a pair of lateral horns. We hypothesize that the initial expression of *HOXD1* in anterior crest cells made this evolutionary co-option possible and thus helped to determine the position and number of horns, which became the distinctive trait of Bovidae.

## Materials and Methods

### Ethics Statement

All experiments reported in this work comply with the ethical guidelines of both the French National Research Institute for Agriculture, Food and Environment (INRAE) and the University of Geneva, Switzerland. Blood samples were collected on sheep and goats during routine blood sampling (for annual prophylaxis, paternity testing or genomic selection purpose) by trained veterinarians and following standard procedures and relevant national guidelines. Sample collection of small ruminants in Switzerland was approved by the Cantonal Committee for Animal Experiments (Canton of Bern; permit 75/16). Ovine and caprine fetuses were produced in an INRAE experimental farm (Bressonvilliers, France) and collected in the INRAE experimental slaughterhouse of Jouy-en-Josas (France). Experiments were performed in strict accordance with the European directive 2010/63/UE and were approved by the local Institutional Animal Care and Use Committee of AgroParisTech/INRAE (COMETHEA, permit number 19/032). All experiments with mice were performed in agreement with the Swiss law on animal protection (LPA), under license No GE 81/14 (to D.D.). All the samples and data analyzed in the present study were obtained with the permission of breeders, breeding organizations and research group providers.

### Animals

Live sheep and goats. Sheep and goats from a wide diversity of breeds around the world were involved in at least one of the analyses performed in this study. Briefly, they felt into four categories: 1) animals genotyped with Illumina OvineHD or GoatSNP50 (Tosser-Klopp et al. 2014) BeadChip for mapping the *POLYCERATE* locus in both species (**Supplementary Tables 1** and **6**); 2) a set of whole genome sequences used for identifying and filtering candidate mutations (**Supplementary Table 13**); 3) animals genotyped by PCR and Sanger sequencing for candidate mutations (**Supplementary Tables 3** and **7**); and 4) polycerate sheep animals genotyped for verifying putative differences between heterozygous and homozygous individuals in terms of distances between the lateral horns and between the upper horns (**Supplementary Table 12**).

Mouse models. Five different transgenic mouse stocks were used (see **Supplementary Table 14**). The *HoxD^(Del365)^* allele was produced by CRISPR-Cas9 technology. sgRNA were designed manually, ordered as DNA oligos at Eurogentec and cloned into px330. sgRNAs were synthetized with HiScribe T7 high yield RNA synthesis kit (New England Biolabs), incubated together with Cas9 mRNA and then electroporated into fertilized mouse zygotes (see also **Supplementary Note 1**). *The HoxD^(Del151)^* allele was obtained by using CRE-mediated recombination (Andrey et al. 2013). The Transgenic fetuses from four strains containing different lacZ constructions were collected from stage E12.5 to E.15.5. The *Hoxd1^Lac^* strain was obtained by inserting a LacZ cassette in the HindIII site of the second exon of *Hoxd1* (Zakany et al. 2001). The BAC^*HoxD*^ and *BAC^Mtx2^* result from the introduction of a LacZ-SV40promoter-*Hoxd1*-zeocin cassette in the HindIII site of the second exon of *Hoxd1* (Schep et al. 2016). These BACs were selected based on their localization on the physical map of the mouse genome (Gregory et al. 2002) and obtained from the RPCI-23 and -24 Mouse (C57BL/6J) BAC Libraries from the Children’s Hospital Oakland Research Institute (https://bacpacresources.org/libraries.php). The modified BACs were purified, linearised and microinjected into mouse fertilised oocytes to obtain each of these strains in mixed Bl6XCBA hybrid background, by standard procedures. Gene expression analyses were performed on heterozygous specimens.

A precise map of the orthologous *Hoxd* region in mouse and goat was obtained by aligning on murine GRCM38/mm10 genome assembly the BAC end sequences and goat genome sequences of 10 kb segments encompassing the breakpoints of the large insertion-deletion. Alignments were carried out using the BLAT tool from the UCSC Genome Browser (http://genome.ucsc.edu/cgi-bin/hgBlat).

Animals subject to post-mortem clinical examination. The eyelids and eyes fundus were examined in a 3-weeks old polycerate male Provençale kid who died from a natural cause and a matched control, as well as in an 8-year old polycerate Jacob ewe and her wild type half-sister after slaughter.

Ovine and Caprine fetuses were generated by mating heterozygous polycerate males of the caprine Provençale and ovine Jacob breeds with wild type cull females, after oestrus synchronization. Oestrus cycles were synchronized using intravaginal sponge impregnated with progestagen for 15 days followed by PMSG (Pregnant Mare Serum Gonadotropin) injection 48 h after sponge removal. Pregnant females were anaesthetized by electronarcosis and euthanized by immediate exsanguination on day 70 or 76 *post-coïtum* in the INRAE slaughterhouse of Jouy-en-Josas (France). Directly after, the fetuses were recovered from their genital tracts and exsanguinated. ‘Skin’ samples comprising the epidermis, dermis and hypodermis were collected at different locations on the left side of the head of the 70 dpc goat and 76 dpc sheep fetuses (see **Fig. 3**) for expression studies. Of note, the skin of the back of the head was sampled slightly more caudally in polycerate animals due to the specific localization of the posterior pair of horns. The same skin samples were collected on the right side of the head with the underlying bone for histological analyses. Four case fetuses and four controls of matched sex were selected in each species for expression studies. Finally, for verification, liver samples were also collected for DNA extraction and subsequent genotyping of the fetuses for the sheep and goat *POLYCERATE* mutations.

Skull specimens. The skulls from 61 sheep (32 polycerate, 29 wild type) and 19 goats (12 polycerate, 7 wild type) were obtained from different anatomical collections. These specimens were sampled over the last 170 years and originate from a wide variety of populations. Information on horn phenotype, species, gender, age, population or breed, collection, and year of entry in the collection are presented in **Supplementary Table 10**.

### Phenotyping

The polycerate phenotype is an autosomal dominant trait readily visible on fetuses at 70 dpc in goat and 76 dpc in sheep (**Supplementary Fig. 8**). Phenotyping at birth is difficult due to the presence of hairs and it is necessary to wait for after the first month to distinguish horns growing amid fur. In polycerate animals, horns have a nearly circular cross section but, depending on their relative placement, they may progressively fuse at the base with other horns located on the same side of the skull. The growth in width of horns is expected to affect the measure of distances between the lateral horns (dlh) and the upper horns (duh) but not their relative sizes. This, together with the fact that we never observed any case of fusion between the upper horns led us to consider the dlh/duh ratio on the left side of the head to distinguish different types of four-horned animals in one of the analyses performed in this study. Polyceraty is frequently associated with defects of the eyelid in both species. While we did not systematically record this particular phenotype, we performed *post-mortem* clinical examination of the eyelids and eyebrows in one case and one control animal per species (**Supplementary Fig. 5-7**).

### DNA Extraction

Ovine and caprine DNAs were extracted from hair root, blood or liver samples using the DNeasy Blood and Tissue Kit (Qiagen). Murine DNA was isolated from ear snip after Proteinase K digestion using standard phenol/chloroform protocol. DNA quality was controlled by electrophoresis and quantified using a Nanodrop spectrophotometer (Thermo Scientific).

### IBD-Mapping of Caprine and Ovine *POLYCERATE* Loci

General principle. Assuming autosomal dominant inheritance and genetic homogeneity in each of the species investigated, all polycerate animals share at least one copy of the same causative mutation and of a surrounding chromosomal segment inherited-by-descent from a common ancestor. Therefore, comparing SNP array genotyping data of two distantly related polycerate animals is expected to reveal a number of Mendelian incompatibilities (i.e. homozygosity for different alleles) throughout their genomes but not within shared IBD segments. Accordingly, we screened Mendelian incompatibilities in all the possible pair combinations of polycerate x polycerate (4H4H pairs) and polycerate x wild type (4H2H) individuals. Pairs with a proportion of Medelian incompatibilities below 1 percent of the total number of markers tested were declared as constituted of parent and offspring and were not considered in the analysis. Then, for sliding windows of n markers (n set to 10 in goat and 50 in sheep considering differences in marker density) we scored the numbers of 4H4H pairs and 4H2H pairs for which ‘no’ versus ‘at least one’ Mendelian inconsistency has been recorded. Finally, we compared the contingency tables produced using Fisher’s exact test.

SNP array genotypes, sample and variant pruning. Illumina GoatSNP50 BeadChip genotypes specifically generated for this research and Illumina OvineHD Beadchip genotyping data generated by two previous studies ^8,10^ were considered in the analyses. Polled (i.e hornless) animals were removed from the sheep dataset. Markers with a minor allele frequency below 5% or which were called in less than 95 % of the samples were eliminated. Moreover, in sheep, genotyping data were extracted for markers located in a 10 Mb region (Chr2:127,500,001-138,500,000) corresponding approximately to the *HOXD* gene cluster +/- 5 Mb and encompassing all the mapping intervals of the *POLYCERATE* locus reported in the literature (Greyvenstein et al. 2016; He et al. 2016; Kijas et al. 2016; Ren et al. 2016). The final datasets contained 111 cases, 87 controls and 2’232 markers in sheep and 35 cases, 51 controls and 48’345 markers in goat.

### Analysis of Whole-Genome Sequences

#### Whole-genome sequences

The genomes of one polycerate Provençale goat and one polycerate Jacob sheep were sequenced specifically for this study. Both were born from polycerate X wild type crosses and thus were predicted to be heterozygous for the caprine and ovine causative variants, respectively. Paired-end libraries with a 450 bp (goat) and 235 bp (sheep) insert size were generated using the NEXTflex PCR-Free DNA Sequencing Kit (Biooscientific). Libraries were quantified with the KAPA Library Quantification Kit (Cliniscience), controlled on a High Sensitivity DNA Chip (Agilent) and sequenced on a HiSeq 2500 (with 2*100 bp read length in goat) and a HiSeq 3000 (with 2*150 bp read length in sheep). The average sequence coverage was 16.7 and 11.1 x, for the polycerate goat and sheep individuals, respectively. Additional whole-genome sequences available in public databases were also considered in the analyses. These consisted of FASTQ files (for 10 additional case and 341 control sheep) and of VCF files (for 1160 goat and 838 sheep control individuals) generated by previous studies (see **Supplementary Table 13**). When necessary, the NCBI Genome Remapping Service (https://www.ncbi.nlm.nih.gov/genome/tools/remap) was used to convert positions in VCF files between older and most recent versions of genome assemblies.

#### Read alignment, variant calling and filtering for candidate variants

The sequence reads from FASTQ files were mapped on goat ARS1 (https://www.ncbi.nlm.nih.gov/assembly/GCF_0017_04415.1/) and sheep Oar_v4.0 (https://www.ncbi.nlm.nih.gov/assembly/GCF_000298735.2) genome assemblies using the BWA-MEM software v 0.7.17 with default parameters (Li and Durbin 2009) and converted to bam format with v 1.8 of SAMtools (Li et al. 2009). Duplicate reads were marked using Picard tools v 2.18.2 MarkDuplicates option (http://broadinstitute.github.io/picard) and base quality recalibration and indel realignments were done with v 3.7 of GATK (McKenna et al. 2010). Reads located in the mapping intervals of the ovine and caprine *POLYCERATE* loci +/- 1 Mb were extracted using SAMtools view option before processing to the calling of SNPs and small indels with GATK-HaplotypeCaller in ERC mode. The minimum read mapping quality and phred-scaled confidence threshold were set to 30 for each sample (‘-stand_call_conf 30.0 -mmq 30 -ERC GVCF -variant_index_type LINEAR -variant_index_parameter 128000’). In goats we retained only heterozygous variants found in the heterozygous polycerate individual and absent from 1160 control animals, while in sheep we focused our attention on variants which were shared (either in heterozygous or homozygous state) in all the 11 polycerate sheep (1 Jacob and 10 Sishui Fur Sheep) and absent from the 1179 control animals. Finally, to ensure that we did not miss any candidate variants, we performed a detection of structural variants in the same regions using Pindel (Ye et al. 2009) and a visual examination of the whole genome sequences for 11 goats (1 case, 10 controls) and 22 sheep (11 cases and 11 controls) using IGV (Thorvaldsdóttir et al. 2013). The count command in IGVtools was used to produce ‘.tdf’ files and identify changes in read coverage in the intervals investigated (with parameters: zoom levels = 10, window function = mean, window size = 1000, and extension factor = 500).

### Definition of the Boundaries of the 503 kb Deletion-137 kb Insertion in Goat

The boundaries of variant g.115,652,290_116,155,699delins137kb were reconstructed manually using split read and paired-end read information obtained from IGV. Sequences of reads affected by the mutation were extracted from the .bam file using linux command lines and aligned manually to reconstruct the nucleotide sequence at each fusion point. For verification, amplicons encompassing these fusion points were PCR amplified in a Mastercycler pro thermocycler (Eppendorf) using Go-Taq Flexi DNA Polymerase (Promega), according to the manufacturer’s instructions and primers listed in **Supplementary Table 15**. Amplicons were purified and bidirectionally sequenced by Eurofins MWG (Hilden, Germany) using conventional Sanger sequencing.

### Genotyping of DNA Sequence Variants

SNP and small Indels were genotyped using PCR and Sanger sequencing as described above. PCR primers were designed with Primer3 software (Rozen and Skaletsky 1999) and variants were detected using NovoSNP software (Weckx et al. 2005). Transgene insertions and large insertion-deletion were genotyped by PCR and electrophoresis on a 2% agarose gel. Ovine variant g.132,832,249_132,832,252del was genotyped with primers TTTGGGGCCACACTAGAATC and CCTAGAGGGGGCCTACGAG while caprine and murine variants were genotyped with the primers listed in **Supplementary Table 7** and **14** respectively.

### Analysis of Nucleotide Sequence Conservation at the *HOXD1* Exon 1–Intron-1 Junction

Nucleotide sequences of the *HOXD1* gene in 103 sarcopterygian and tetrapod species were obtained from the Ensembl (http://www.ensembl.org/index.html; release 98) and UCSC (http://genome.ucsc.edu/) genome browser databases. The localization of the nucleotide sequence (between *MTX2* and *HOXD3*) was verified in each genome assembly to avoid possible confusion with paralogs. In addition, only one sequence was arbitrarily retained when genome assemblies for distinct individuals of the same species were available. Then sequences were put in the same orientation and trimmed to get 40 nucleotides before and 20 nucleotides after the splice donor site of *HOXD1* exon 1. A multispecies alignment was generated with ClustalW software (Thompson et al. 1994), version 2.1 (https://www.genome.jp/tools-bin/clustalw) and a sequence logo was generated using WebLogo (Crooks 2004) (http://weblogo.berkeley.edu/). Information on species, sequence and genome assemblies are presented in **Supplementary Table 5**.

### Fluorescence *In Situ* Hybridization in Goat

Skin biopsies were sampled from one heterozygous polycerate and one wild-type fetuses. Fibroblast cultures and metaphases were obtained according to (Ducos et al. 2000). Nucleotide sequences from the segments of caprine chromosomes 2 and 5 involved in the candidate causative mutation were aligned against bovine bacterial artificial chromosome (BAC) end sequences using BLAST (http://blast.ncbi.nlm.nih.gov/Blast.cgi). Two INRA BAC clones (Eggen et al. 2001) were selected and obtained from the Biological Resources of @BRIDGe facilities (abridge.inrae.fr): INRAb 230B11, targeting the segment deleted on Chr2, and INRAb 348A12, targeting the region of Chr5 that is duplicated and inserted on Chr2. FISH experiments were carried out according to (Yerle et al. 1994). The two BACs were labeled with biotin and digoxygenin, respectively, using the BioPrime DNA Labeling System kit (Life Technologies, Carlsbad, CA, USA). Finally, they were revealed by Alexa 594 conjugated to streptavidin (Molecular Probes, Eugene, OR, USA) and FITC conjugated mouse anti-digoxygenin antibodies (Sigma, St Louis, MO).

### Histological Analyses

Tissues were fixed in paraformaldehyde (4%) for 24 h at +4°C. Samples were subsequently dehydrated in a graded ethanol series, cleared with xylene and embedded in paraffin wax. Microtome sections (5 μm, Leica RM2245) were mounted on adhesive slides (Klinipath-KP-PRINTER ADHESIVES), deparaffinized, and stained with haematoxylin, eosin and saffron (HES). Slides were scanned with the Pannoramic Scan 150 (3D Histech) and analyzed with the CaseCenter 2.9 viewer (3D Histech).

### Quantitative RT-PCR

RNA was extracted using the RNeasy Mini Kit (Qiagen). Super-Script II (Invitrogen) was used to synthesize cDNA from 2 μg of total RNA isolated from each tissue sampled in 70 dpc goat and 76 dpc sheep fetuses. Gene sequences were obtained from Ensembl v92 (www.ensembl.org) and PCR primers (**Supplementary Table 16**) were designed using Primer Express Software for Real-Time PCR 3.0 (Applied Biosystems). Primer efficiency and specificity were evaluated on genomic DNA in each species. Quantitative PCR was performed in triplicate with 2 ng of cDNA using the Absolute Blue SYBR Green ROX mix (Thermo Fisher Scientific) and the StepOnePlus Real-Time PCR System (Applied Biosystems). The expression stability of five genes (*RPLP0, GAPDH, H2AFZ, YWHAZ* and *HPRT1*) was tested at each time point using the GeNorm program (Vandesompele et al. 2002) to identify appropriate qRT-PCR normalizing genes. Three normalizing genes (*GAPDH, H2AFZ* and *HPRT1*) were retained and the results were analyzed with qBase software (Hellemans et al. 2007).

### Evaluation of the Consequences of Intron Retention Due to the 4 bp Deletion in *HOXD1* intron 1

The complete nucleotide sequence of ovine *HOXD1* gene was obtained from Ensembl v97. A mutant mRNA characterized by (i) a retention of intron 1 and (ii) a deletion of nucleotides located at position +4 to +7 bp after the end of exon 1 was designed. This mutant mRNA was translated using ExPASy Translate tool (https://web.expasy.org/translate/). Information on HOXD1 functional domains was obtained from UniProt Knowledgebase (https://www.uniprot.org/uniprot/W5Q7P8).

### 3D Geometric Morphometrics

#### Three-dimensional models

Three-dimensional models were generated for 80 skulls consisting of 32 polycerate and 29 wild type sheep specimens as well as 12 polycerate and 7 wild type goat specimens (for information on skulls and reconstruction methods see **Supplementary Table 10**). Most of the 3D models (n=47) were reconstructed using a Breuckmann StereoScan structured light scanner and its dedicated software OptoCat (AICON 3D systems, Meersburg, Germany). Twenty-nine skulls were digitized with the Artec Eva structured-light scanner and ScanStudioHD software v12.1.1.12 (Artec 3D, Luxembourg, Luxembourg). In addition, four skulls were digitized with a photogrammetric approach, similar to that described in (Evin et al. 2016). In brief, hundred pictures per sample were taken on different angles and inclinations with a Nikon D3300 camera equipped with an AF-S Micro Nikkor 85mm lens (Nikon, Tokyo, Japan) and a self-made fully automatic turntable. Then 3D models were reconstructed with the ReCap Photo software (Autodesk, San Rafael, CA, USA). Previous studies indicated no significant differences between 3D models obtained with three-dimensional scanners or photogrammetry (Evin et al. 2016; Fau et al. 2016). Both approaches are comparable in terms of measurement error (less than 1 mm). Bone surfaces were extracted as meshes and geometric inconsistencies (i.e. noise, holes) were cleaned using Geomagic software (3D Systems, Rock Hill, USA).

#### Shape analyses

116 3D landmarks and sliding semi-landmarks were placed on each specimen by the same operator using the IDAV Landmark software (Wiley et al. 2005) v3.0. Out of them 16 were anatomical landmarks, and 100 were sliding semi-landmarks individually placed around the basis of the horns on the suture between the bony core and the frontal bone. On each side, the first of these 50 sliding semi-landmarks was placed on the upper horn, at the intersection between the upper ridge of the bony core and the suture previously mentioned. Details on landmark locations on polycerate and wild type specimens are provided in **Supplementary Table 11** and **Supplementary Fig. 11**.

Following the procedure detailed by (Botton-Divet et al. 2015), a template was created using the specimen 2000-438 on which all anatomical landmarks and surface sliding semi-landmarks were placed. Then, a semi-automatic point placement was performed (Gunz and Mitteroecker 2013) to project sliding semi-landmarks on the surface of the other 3D digitized skulls. Sliding semi-landmarks on surfaces and curves were allowed to slide in order to minimize the bending energy of a thin plate spline (TPS) between each 3D meshes and the template. After this first TPS relaxation using the template, three iterative relaxations were performed using the Procrustes consensus of the previous step as a reference.

To remove non-shape variation (i.e. differences in position, scale, and orientation of the configurations) and provide optimal comparability between the specimens, we performed a generalized Procrustes Analysis (GPA) (Rohlf and Slice 1990). Since our dataset contained more variables than observations, we performed a Principal Component Analysis (PCA) on the procrustes residuals to reduce dimensionality, as recommended by (Gunz and Mitteroecker 2013), and plotted the first Principal Components (PCs) to visualize the specimen distribution in the morphospace. In addition, the mean shape of our sample was used to compute theoretical shapes associated with the maximum and minimum of both sides of the first PC axis for each species using thin plate spline. GPA, PCA and shape computations were done using the ‘Morpho’ and ‘geomorph’ packages (Adams and Otárola-Castillo 2013; Adams et al. 2018; Schlager 2018) in the R environment (R Core Team 2018).

#### Repeatability and reproducibility of landmark placement

The 116 landmarks and sliding semi-landmarks were placed ten times independently on the skulls from two polycerate and two control male sheep sampled between 1852 and 1909 in Tunisia (A-12130, A12132, 1909-4) and neighboring Algeria (A12157; see **Supplementary Table 10**). The measurements were superimposed using a GPA and analyzed using a PCA. Since the variation within specimens was clearly smaller than the variation between specimens (**Supplementary Fig. 12**), we considered that the 116 landmarks and sliding semi-landmarks were precise enough to describe shape variation.

## Data Availability

Raw sequencing data that support the findings of this study have been deposited to the European Variation Archive (EVA, https://www.ebi.ac.uk/eva/) under accession number PRJEB39341. Sequences from previous studies can be found at the following URL (www.goatgenome.org/vargoats_data_access.htm) or in the NCBI BioProject and EVA databases under accession numbers PRJEB6025, PRJEB6495, PRJEB9911, PRJEB14098, PRJEB14418, PRJEB15642, PRJEB23437, PRJEB31241, PRJEB31930, PRJEB32110, PRJEB35553, PRJEB35682, PRJEB37460, PRJEB39341, PRJEB39341 and PRJNA624020. Illumina GoatSNP50 Beadchip genotyping data generated for this study have been deposited in the Dryad Digital Repository (doi: 10.5061/dryad.rxwdbrv6n). Illumina OvineHD Beadchip genotyping data from previous studies can be found in the same repository (doi: 10.5061/dryad.6t34b and 10.5061/dryad.1p7sf). Coordinates of landmarks and source data underlying Fig. 3 and 4, and Suppl. Fig. 2, 10 and 11 are provided as a Source Data file.

## Acknowledgements

We thank L. Orlando (Universities of Toulouse III, France and Copenhagen, Denmark) for tentatively extracting DNA from museum skull specimens, as well as C. Hozé, F. Lejuste and A Michenet (ALLICE), R. McCulloch (CSIRO), S. Chahory and C. Degueurce (ENVA), F. Andreoletti, M. Boussaha, J. Kergosien, D. Mauchand, M. Femenia, N. Perrot and M. Vilotte (INRAE), B. Camus-Allanic (LABOGENA DNA), J. Peters (LMU), C. Bens, A. Delapré and A.Verguin (MNHN), L. Ludes-Fraulob (MZS) and B. Mascrez (University of Geneva) for their assistance. We also thank the staff of the INRAE experimental unit UE 1298 SAAJ for animal husbandry and management, as well as breeders and zoological parcs for making animals available for sampling and for providing pictures. Contributors include in particular the Capgènes breeding company (France), L. Pachot (Mouton Village, Vasles, France), A. Archiloque (France), L. Fiorenzi (Az. Agr. Madonna delle Alpi, Italy), the Schafzuchtverein Jakobschaf Schweiz (Switzerland), Tierpark Hamm (Germany), A. Schumann (Germany) and Dr. A. Ennaifer (Zoological Park of Tunis, Tunisia). Finally, the authors thank the VarGoats Consortium for allowing variant filtering against their dataset. The Vargoats project was supported by France Génomique (grant number ANR-10-INBS-0009). This work was supported by APIS-GENE (grant AKELOS) and the Swiss National Research Foundation (grant number No 310030B_138662 to D.D.). A.Hi. was supported by a PhD fellowship from the University of Geneva.

## Authors contributions

O.G., D.R., T.H., N.C., C.M.R., B.J.H and J.K. provided Illumina OvineHD Beadchip genotyping data and related phenotypes. A.C. mapped the ovine and caprine polycerate loci. C. Dr., C.D.-B., D.B., I.M, L.P., O.G., T.H., G.B., F.M., N.H., J.P., S.B.J., J.H., R.R., I.P., J.A.L., L.G., D.R., E.V.M.-K., N.C., B.J.H, J.K. and G.T.-K. provided samples and phenotypes. D. E. and C.Do. performed whole genome sequencing from one polycerate Provençale goat and one polycerate Jacob sheep. G.T.-K. provided control whole genome sequences from sheep and goats. A.C., P.B., and M.N.-S. performed variant calling, annotation and screening for candidate variants. A.C. and A.A.-B. analysed sequence conservation and annotated the gene content of the mapping intervals. M.-C.D., C.Gr. and A.Hi. extracted DNA. M.-C.D., A.C., C.Gr., and A.Hi. performed PCR for Sanger sequencing and for genotyping by PCR and electrophoresis or PCR and Sanger sequencing. A.P. performed FISH analysis. A.Hi., J.Z and D.D. produced and studied mouse models. E.P. provided access to laboratory and experimental farm facilities. A.C., A.A.-B., M.-C.D., C.Gr. and E.P. sampled ovine and caprine fetuses. A.A.-B. and M.-C.D. extracted RNA, performed qRT-PCR and analysed the results. A.Bo., J.R., A.C., A.A.-B. and M.-C.D. performed histological analyses. A.C., M.S. and A.Ha. performed 3D data acquisition of skulls. A.Bl. provided access to a light scanner for 3D data acquisition. O.P., J.L., R.S., M.-D.W., R.-M.A. and C.Gu. provided access to skull specimens and related information. A.C. performed morphometric analyses. R.C. provided software and expertise in morphometric analyses. A.C. (Bovidae) and D.D. (mouse) designed the studies and wrote the manuscript, which was accepted or revised by all authors.

