## Supplemental figures, tables and note for "Analysis of Polycerate Mutants Reveals the Evolutionary Co-option of *HOXD1* to Determine the Number and Topology of Horns in Bovidae"

This document contains:

Supplementary Figures 1-13,

Supplementary Tables 1-16,

Supplementary Note 1,

and additional references.

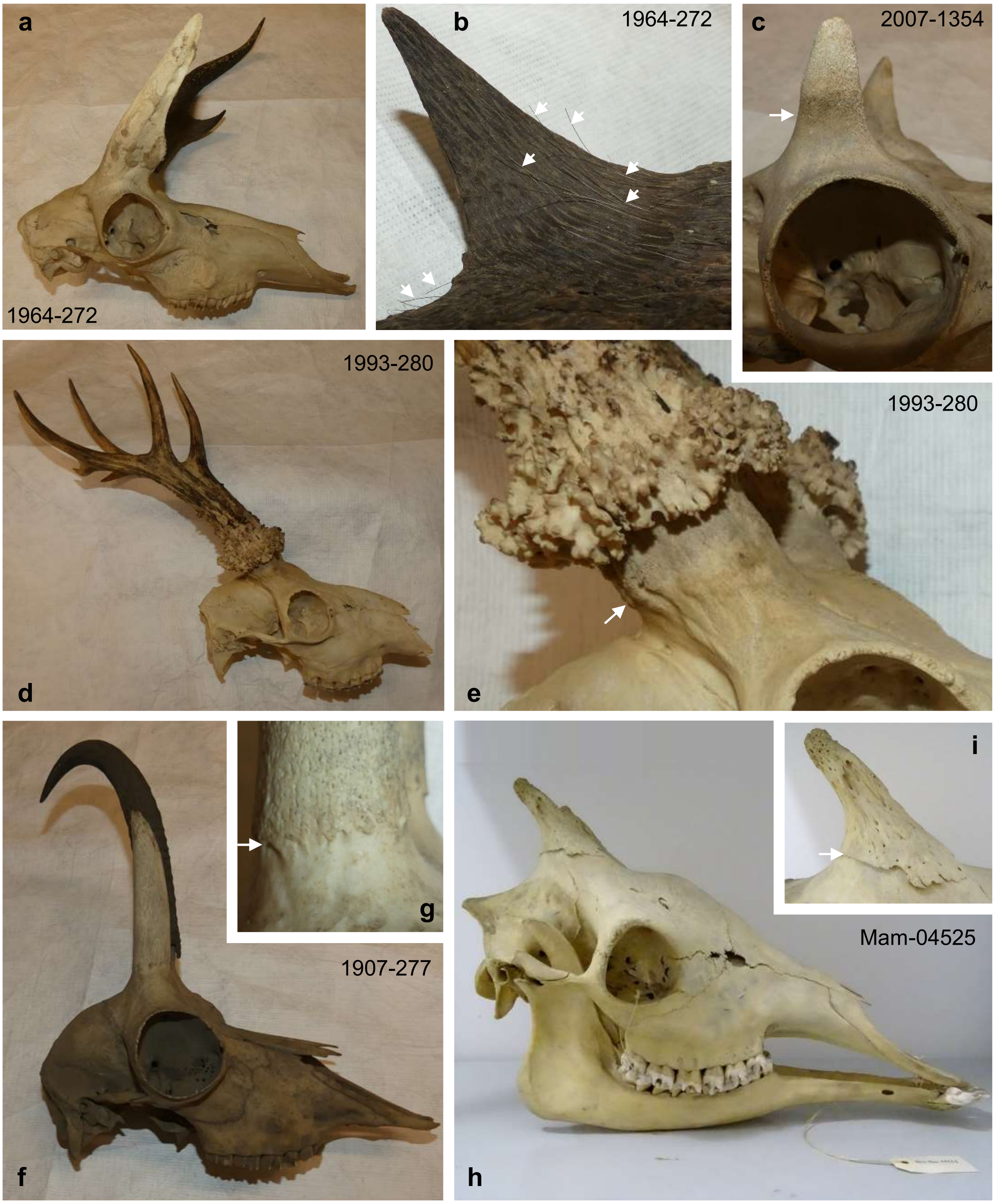


**Supplementary Figure 1. Comparison of Pecoran headgear types.** a-c) Pronghorns of *Antilocapra Americana* in adult male (a,b) and female (c) skull specimens. d,e) Antlers of a male *Capreolus capreolus*. f,g) Horns of a male *Rupicapra rupicapra*. h,i) Ossicones of a *Giraffa Camelopardalis* of unknown gender. Note the shared position of the different headgear types on the frontal bones (a, d, f, h) and the change of texture between the frontal bone and their bony core as indicated by arrows (c, e, g, i). Note also the presence of vestigial hairs (marked with arrows) after keratinization of the decidual sheath of the pronghorn in (b). Specimens MNHN-ZM-AC 1964-272, 2007-1354, 1993-280 and 1907-277 belong to the *Collection d'anatomie comparée du Muséum National d'Histoire Naturelle*, Paris, France while Mam-04525 originates from the *Collection d'anatomie du Musée Zoologique de Strasbourg*, France.

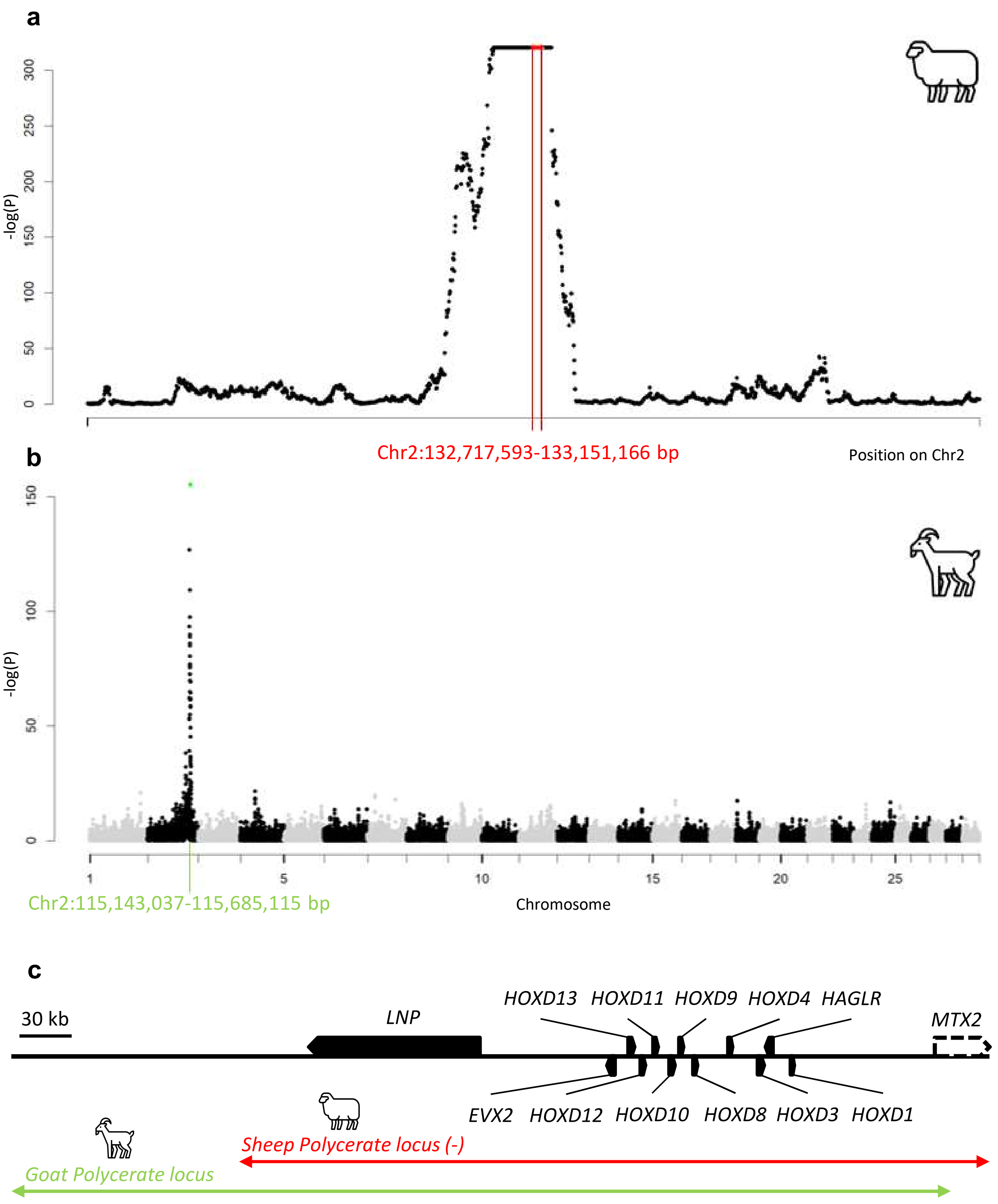


**Supplementary Figure 2. Mapping of the *POLYCERATE* loci in sheep and goat.** a) Identical-By-Descent (IBD) mapping of ovine *POLYCERATE* locus around the *HOXD* gene cluster and +/- 5 Mb on Chr2 (see Methods). Fisher's exact test p-values equal to zero were set to 1.0 x $10^{-320}$; 30 consecutive sliding windows of 50 markers corresponding to an IBD segment shared by all the polycerate sheep are highlighted in red. b) IBD mapping of caprine *POLYCERATE* locus on Chr2. One IBD segment of 10 markers shared by all the polycerate goats is highlighted in green. The intervals indicated are defined by the positions of the most proximal markers outside of the IBD segments (on Oar_v4.0 and ARS1 assemblies). c) Gene content and relative localization of the mapping intervals of the *POLYCERATE* loci in sheep (minus strand) and goat. Dashed lines indicate that a part of *MTX2* is located outside of the region displayed. Sheep and goat icons were made by "Monkik" from www.thenounproject.com.

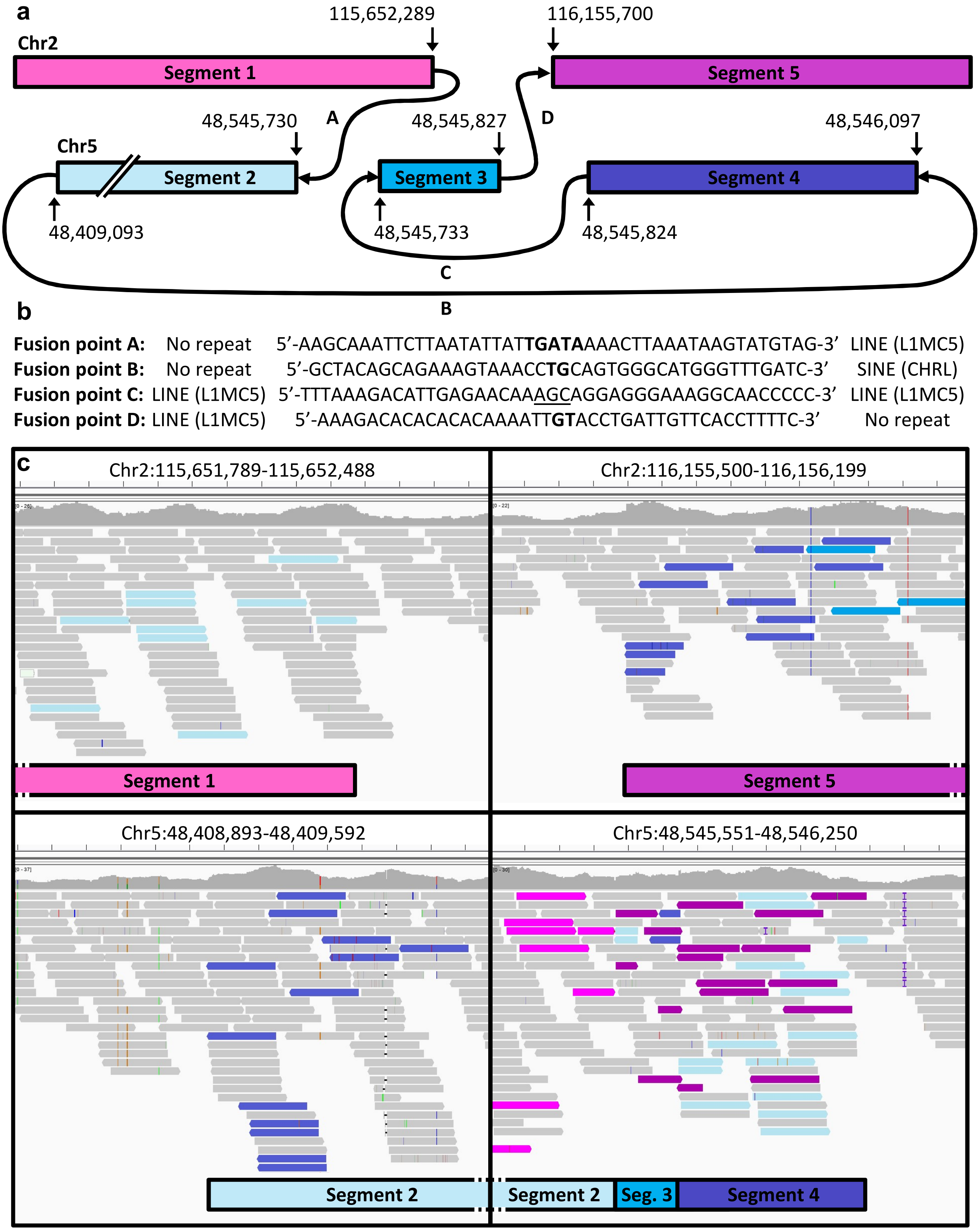


**Supplementary Figure 3. Details on the large insertion-deletion at the goat *POLYCERATE* locus.** a) Schematic representation of the segments involved in the translocation of 137 kb from Chr5 and the 503-kb deletion on Chr2. Fusion points are coded with letters. Positions refer to ARS1 caprine genome assembly. b) DNA sequences at the four fusion points. Nucleotides in bold highlight microhomologies between the fused segments, whereas nucleotides inserted between the breakpoints are underlined. The existence of repeated elements on each side of fusion points is indicated. c) Integrated Genome Viewer screen-captures showing aligned paired-end reads (~100-bp) around the various breakpoints. Reads from aberrantly mapping pairs are color-coded to indicate the chromosomal segment to which the other pair maps.

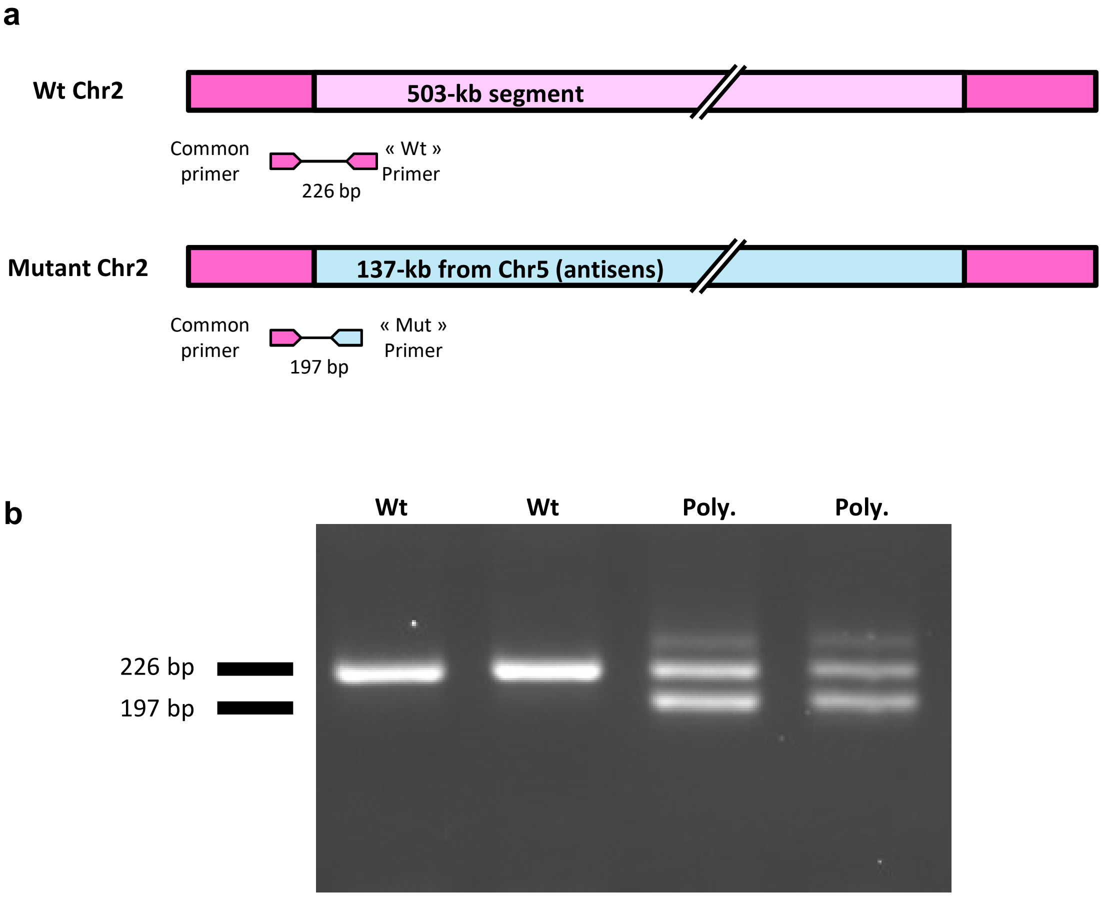


**Supplementary Figure 4. Genotyping of g.115,652,290_116,155,699delins137kb in goat using PCR and electrophoresis.** a) Schematic representation of the segments involved in the mutation and localization of the PCR primers on wild type (Wt) and mutant chromosomes. The coordinates of primers « Common », « Wt », and « Mut » on ARS1 genome assembly are Chr2:115,652,119-115,652,140, Chr2:115,652,323-115,652,344, and Chr5:48,545,700-48,545,722, respectively. b) Agarose gel electrophoresis of amplicons from wild type (Wt) and polycerate (Poly.) animals.

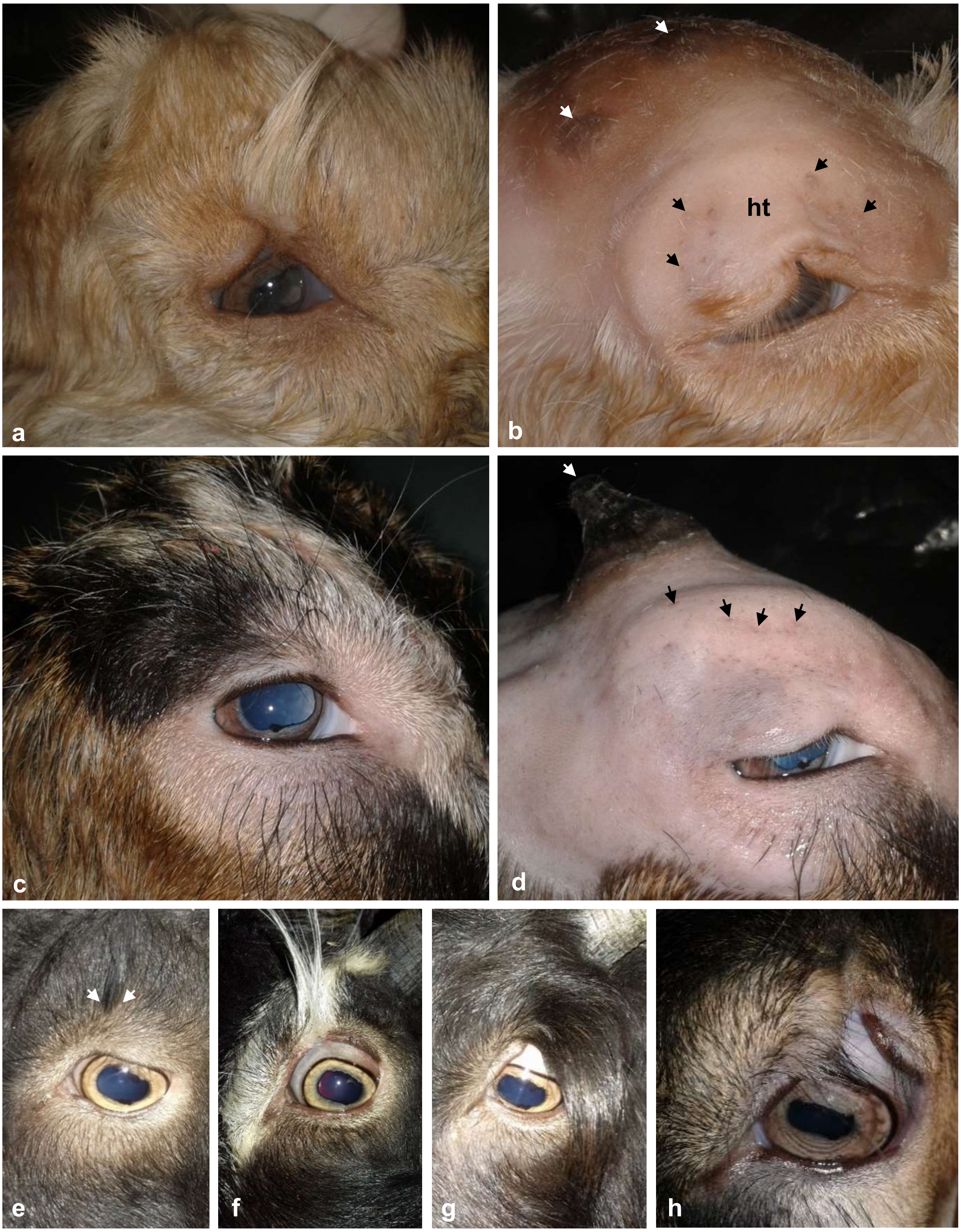


**Supplementary Figure 5. Details on eyelid and eyebrow malformation in polycerate goats.** Upper head of a one-week old polycerate (a, b) and a four-week old wild type (c, d) billy-goat before and after shaving. Note the splitting of the eyelid, of the two rows of vibrissae forming the eyebrow and of the horn buds (white arrows), as well as the presence of abnormally long hair forming a tuft (ht) in the polycerate individual. Black arrows point to vibrissae from the upper row of the eyebrow that are orthologous in (b) and (d) **.** e-h) Eyelid malformation of increasing severity in adult polycerate male goats of the Provençale breed. e) absence of eyelid malformation and presence of a small hair tuft marked with arrows, f) notch in the eyelid margin and hair tuft of moderate size, g) eyelid coloboma partially hidden by a hair tuft of important size. h) Details of the eyelid in (g) after cutting the hair tuft.

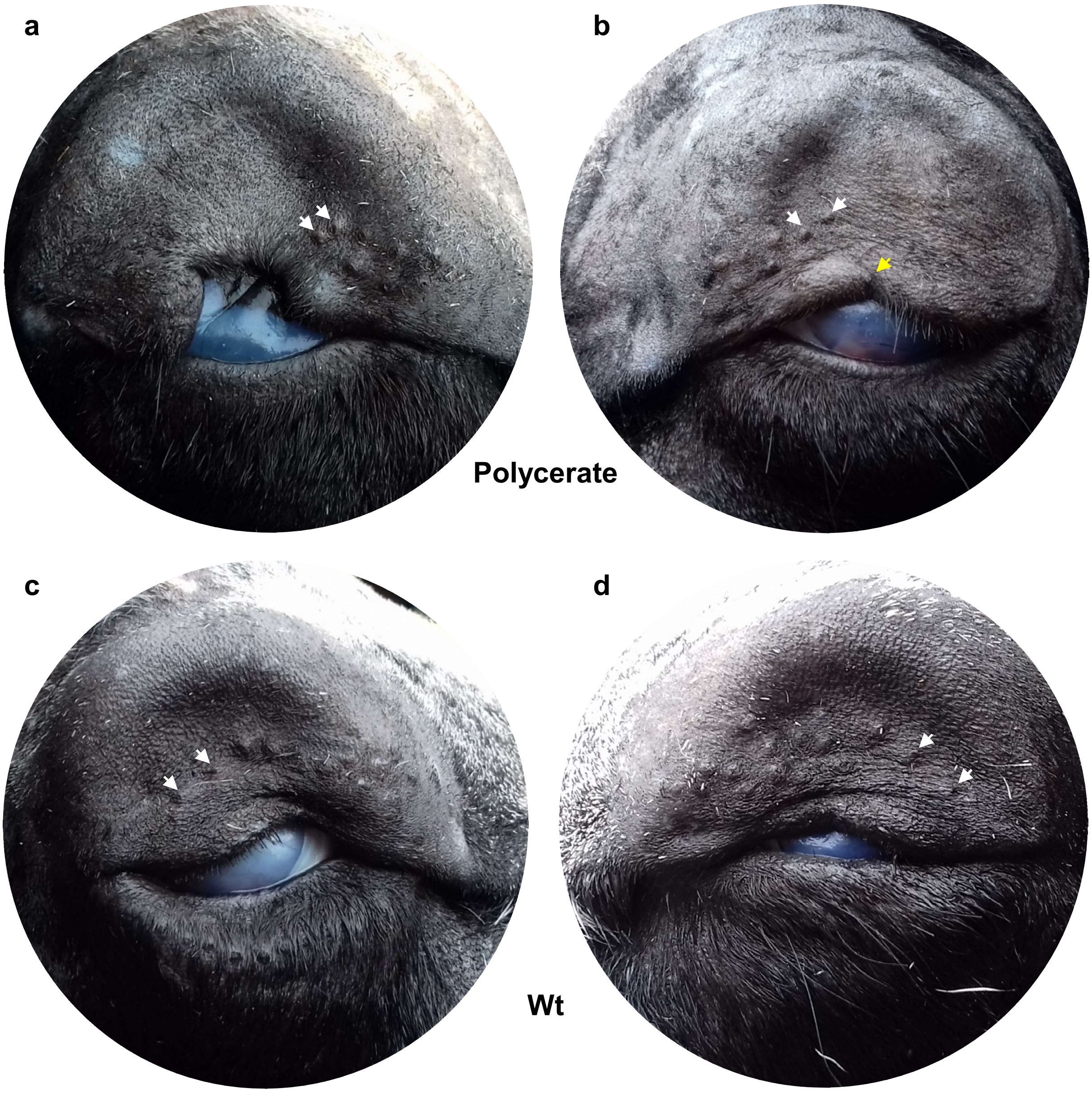


**Supplementary Figure 6. Details on eyelid and eyebrow malformation in polycerate sheep.** Right and Left eyes of a polycerate (a, b, respectively) and a wild type (c, d, respectively) adult Jacob ewe, after shaving. Note the shortened eyebrows in the polycerate ewe as shown by the white arrows pointing to the two most posterior vibrissae. Note also the asymmetry of the eyelid defect with a large portion of the eyelid missing in (a) and only a small notch (yellow arrow) in (b). No hair tuft was observed before shaving. To our knowledge, the presence of abnormally long hair on the eyelid is only observed in polycerate goats. The white color of the cornea is due to the freezing of head samples after slaughter for conservation purpose.

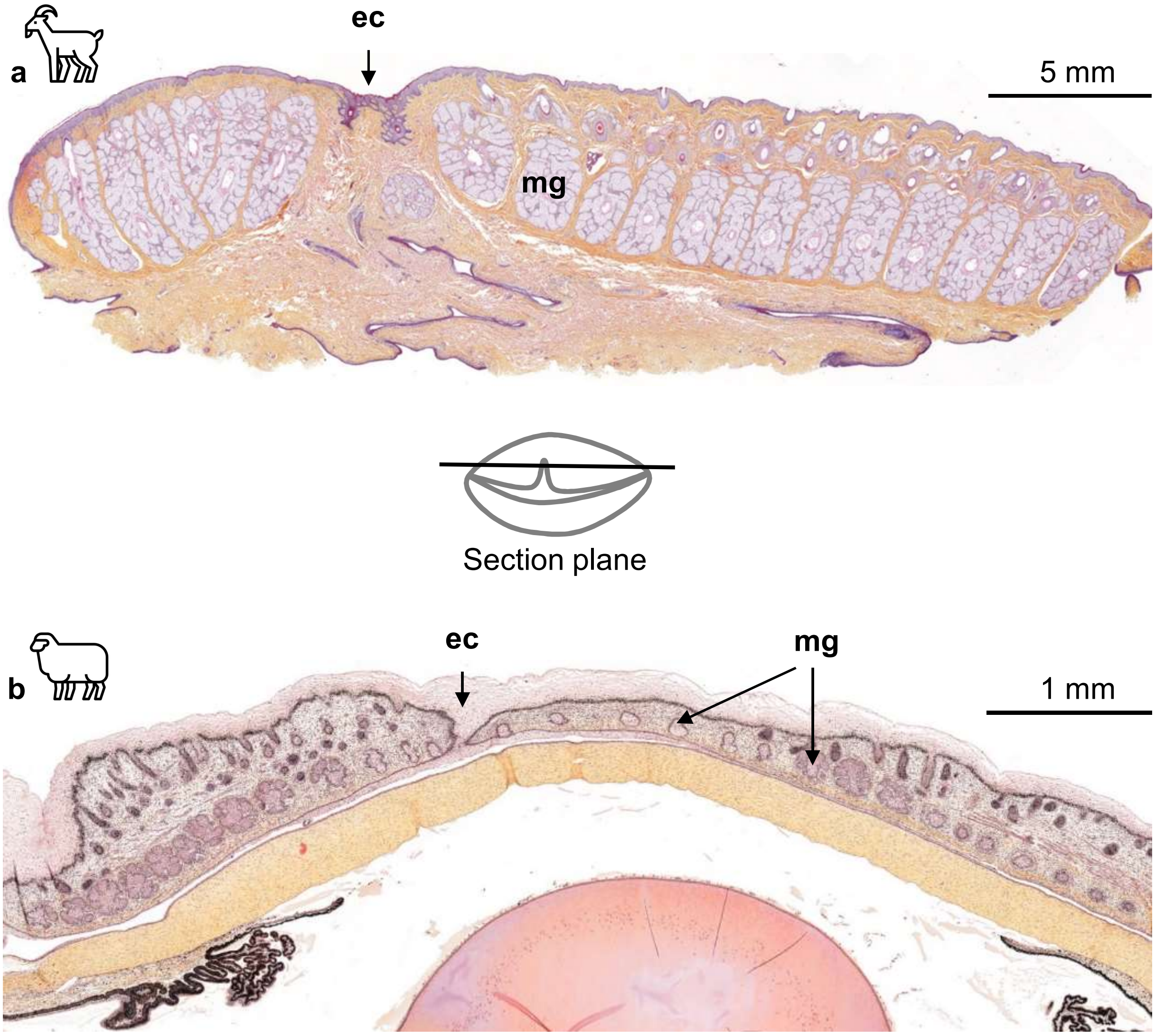


**Supplementary Figure 7. Histology of split upper eyelids in polycerate goats and sheep.** Longitudinal section of the upper eyelid of an adult heterozygous polycerate billy-goat (a) and a heterozygous polycerate female sheep fetus at 76 dpc (b). Meibomian glands (mg) are either absent or underdeveloped in the vicinity of the eyelid coloboma (ec). Sheep and goat icons were made by “Monkik” from www.thenounproject.com.

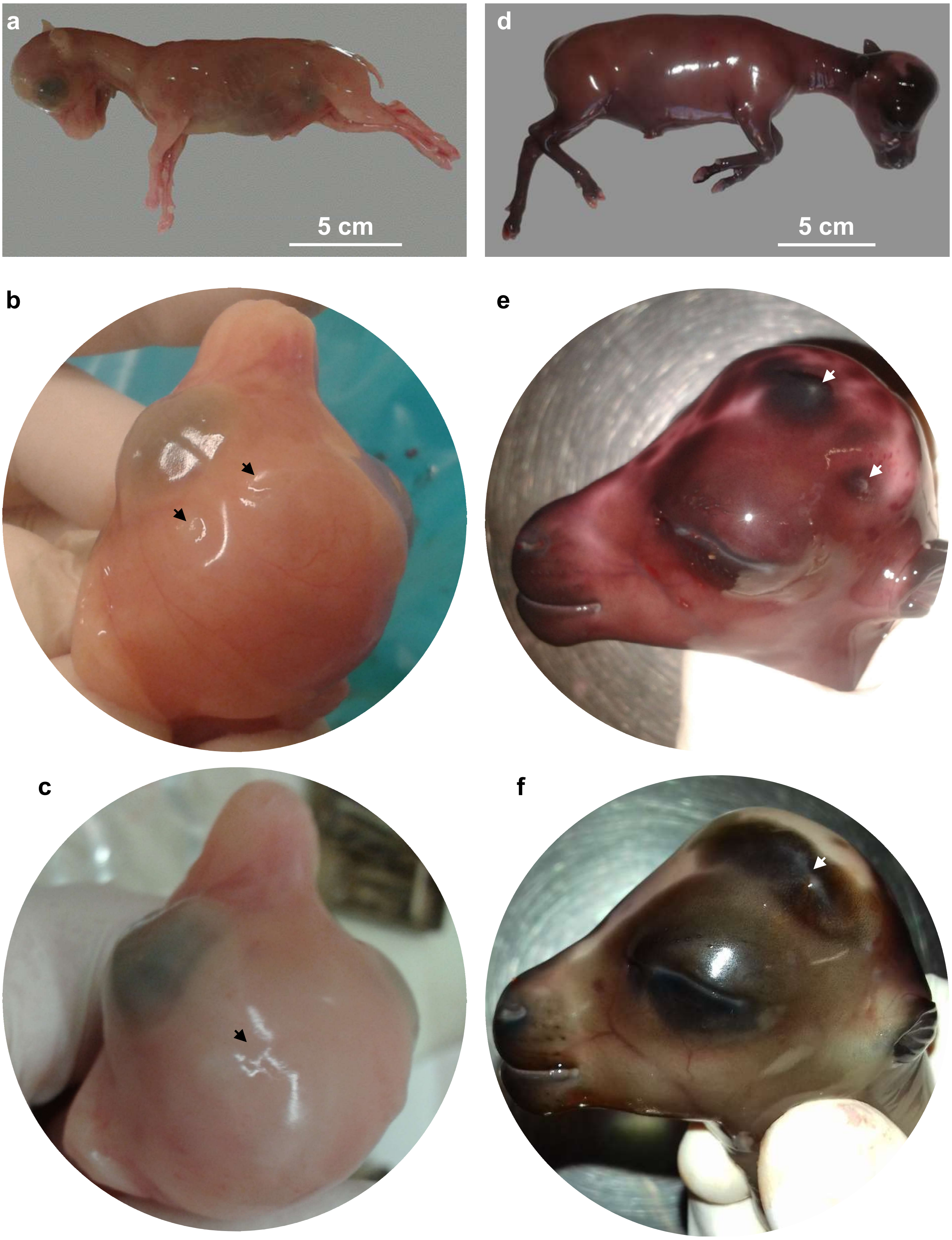


**Supplementary Figure 8. Pictures of goat and sheep fetuses.** a,d) General view of goat (a) and sheep (d) fetuses at 70 and 76 dpc, respectively. b, c) Head of polycerate (b) and wild type (c) goat fetuses. e, f) Head of polycerate (e) and wild type (f) sheep fetuses. Arrows indicate horn buds.

**Supplementary Figure 9. cDNA (top) and amino acid (bottom) sequences for ovine wild type (Wt) and polycerate (Mut) *HOXD1* alleles.** Nucleotide sequences are represented in black for exon 1, red for intron 1, and blue for exon 2. Codons are labelled with alternating yellow triplets. The two nucleotides flanking the 4 bp deletion in the polycerate (Mut) allele are bolded and underlined. Boxes correspond to the RT-qPCR amplicons from intron 1 and exon 2 used for confirming intron retention. Finally, the homeobox DNA binding site of the wild type (Wt) protein is highlighted in green.

```
Wt      1 ATGAGCTCCTACCTCGACTACGTGTCGTGCGGCCGGGACGGCGGCGACTTGCTGAGCTTC    60
Mut     1 ATGAGCTCCTACCTCGACTACGTGTCGTGCGGCCGGGACGGCGGCGACTTGCTGAGCTTC    60
Wt      1 -M--S--S--Y--L--D--Y--V--S--C--G--R--D--G--G--D--L--L--S--F-    20
Mut     1 -M--S--S--Y--L--D--Y--V--S--C--G--R--D--G--G--D--L--L--S--F-    20

Wt     61 GCGCCCAAGTTCTGTCGCGCCGACGCCCGGCCCATGGTCCTGCAGCCCGCCTTCCCCCTG   120
Mut    61 GCGCCCAAGTTCTGTCGCGCCGACGCCCGGCCCATGGTCCTGCAGCCCGCCTTCCCCCTG   120
Wt     21 -A--P--K--F--C--R--A--D--A--R--P--M--V--L--Q--P--A--F--P--L-    40
Mut    21 -A--P--K--F--C--R--A--D--A--R--P--M--V--L--Q--P--A--F--P--L-    40

Wt    121 GGCAGCGGCGACGGCGCGTTTGTTAGCTGCCTGCCCCTGGCCGCGGCGCGAGCGGGGCCC   180
Mut   121 GGCAGCGGCGACGGCGCGTTTGTTAGCTGCCTGCCCCTGGCCGCGGCGCGAGCGGGGCCC   180
Wt     41 -G--S--G--D--G--A--F--V--S--C--L--P--L--A--A--A--R--A--G--P-    60
Mut    41 -G--S--G--D--G--A--F--V--S--C--L--P--L--A--A--A--R--A--G--P-    60

Wt    181 TCGCCCCCGGCCGCCCCGGCGCAGCCTCCCGGGCCTGCCGCGCCCGCGTACGCGCCCTGC   240
Mut   181 TCGCCCCCGGCCGCCCCGGCGCAGCCTCCCGGGCCTGCCGCGCCCGCGTACGCGCCCTGC   240
Wt     61 -S--P--P--A--A--P--A--Q--P--P--G--P--A--A--P--A--Y--A--P--C-    80
Mut    61 -S--P--P--A--A--P--A--Q--P--P--G--P--A--A--P--A--Y--A--P--C-    80

Wt    241 CCCCTAGAGGGGGCCTACGAGCCAGGCGCCGCACCTGTCCAGGCCGGGGGCGAGGACTGC   300
Mut   241 CCCCTAGAGGGGGCCTACGAGCCAGGCGCCGCACCTGTCCAGGCCGGGGGCGAGGACTGC   300
Wt     81 -P--L--E--G--A--Y--E--P--G--A--A--P--V--Q--A--G--G--E--D--C-   100
Mut    81 -P--L--E--G--A--Y--E--P--G--A--A--P--V--Q--A--G--G--E--D--C-   100

Wt    301 TGCCTGCCGGGGTCTGCGCCCGCATACGAGTTACCGTGCGCGCTCGGGCGGCCGGCAGAC   360
Mut   301 TGCCTGCCGGGGTCTGCGCCCGCATACGAGTTACCGTGCGCGCTCGGGCGGCCGGCAGAC   360
Wt    101 -C--L--P--G--S--A--P--A--Y--E--L--P--C--A--L--G--R--P--A--D-   120
Mut   101 -C--L--P--G--S--A--P--A--Y--E--L--P--C--A--L--G--R--P--A--D-   120

Wt    361 GACAGCGGGGCGCACGTCCATTACCCGCCCCCCGCCCCCGGCGTCTCCCCCAAGTGTGCG   420
Mut   361 GACAGCGGGGCGCACGTCCATTACCCGCCCCCCGCCCCCGGCGTCTCCCCCAAGTGTGCG   420
Wt    121 -D--S--G--A--H--V--H--Y--P--P--P--A--P--G--V--S--P--K--C--A-   140
Mut   121 -D--S--G--A--H--V--H--Y--P--P--P--A--P--G--V--S--P--K--C--A-   140

Wt    421 TCCCCAGCCTCCGGCCTCCCTGCCGCCTTCAGCACGTTCGAGTGGATGAAAGTGAAGAGG   480
Mut   421 TCCCCAGCCTCCGGCCTCCCTGCCGCCTTCAGCACGTTCGAGTGGATGAAAGTGAAGAGG   480
Wt    141 -S--P--A--S--G--L--P--A--A--F--S--T--F--E--W--M--K--V--K--R-   160
Mut   141 -S--P--A--S--G--L--P--A--A--F--S--T--F--E--W--M--K--V--K--R-   160

Wt    481 AACGCGCCGAAGAAAAGCAAATTCGCGGAGTATGGAGCCGCCACCCCCTCCAGCGCGATC   540
Mut   481 AACGCGCCGAAGAAAAGTACTTGAGCCTTGGACGGACGGACGCGACTGGGGTTGGGGACG   540
Wt    161 -N--A--P--K--K--S--K--F--A--E--Y--G--A--A--T--P--S--S--A--I-   180
Mut   161 -N--A--P--K--K--S--T--*                                        167

Wt    541 CGCACGAATTTCAGCACCAAGCAACTGACAGAACTGGAGAAGGAGTTTCATTTCAATAAG   600
Mut   541 AGCCTGCCCAGCGGAACTGAGCCGAGGGCGCGCTGCAGCGGTCCCTGTGGACGGCGGTTT   600
Wt    181 -R--T--N--F--S--T--K--Q--L--T--E--L--E--K--E--F--H--F--N--K-   200
Mut

Wt    601 TACTTAACTCGGGCCCGACGCATCGAGATAGCCAACTCATTGCAACTGAATGACACCCAA   660
Mut   601 GGGTCTTATGGGGAACACGCGGCCCCAGGCATTATTAAACCAGGCGAATTGCATTGAGCG   660
Wt    201 -Y--L--T--R--A--R--R--I--E--I--A--N--S--L--Q--L--N--D--T--Q-   220
Mut
```

```
Wt    661 GTCAAAATCTGGTTCCAGAACCGCAGGATGAAACAGAAGAAAAGGGAACGGGAAGGGCTT   720
Mut   661 TGCCCCGAATGCCAGGGGTTGAGAAAATTGCCGGAAGGAGCGCTCCGTGAAGCTTTTCTC   720
Wt    221 -V--K--I--W--F--Q--N--R--R--M--K--Q--K--K--R--E--R--E--G--L-   240
Mut

Wt    721 CTGCCCTCGGCCACCCCCGTGGCTTCCTTCCAGCTTCCCCTCTCAGGACCGAGCCCTGCC   780
Mut   721 TTGGCACAGCTGTGTTCGGAGGCCCAGAAGGGCCGCTCAGTAACTCTGATTCTAGTGTGG   780
Wt    241 -L--P--S--A--T--P--V--A--S--F--Q--L--P--L--S--G--P--S--P--A-   260
Mut

Wt    781 AAGTCTGGCAAGAACCCGGGGAGCCCCTCTCAGGCCCAGGAGCCATCCTGA            831
Mut   781 CCCCAAAGCTCTCCCAAGTCGCGTCTCCGTGTCATCTCGCCACGCTGACGACGCCCTGTC   840
Wt    261 -K--S--G--K--N--P--G--S--P--S--Q--A--Q--E--P--S--*-            276
Mut

Mut   841 TTTACGTTGCAGGCAAATTCGCGGAGTATGGAGCCGCCACCCCCTCCAGCGCGATCCGCA   900
Wt

Mut   901 CGAATTTCAGCACCAAGCAACTGACAGAACTGGAGAAGGAGTTTCATTTCAATAAGTACT   960
Wt

Mut   961 TAACTCGGGCCCGACGCATCGAGATAGCCAACTCATTGCAACTGAATGACACCCAAGTCA  1020
Wt

Mut  1021 AAATCTGGTTCCAGAACCGCAGGATGAAACAGAAGAAAAGGGAACGGGAAGGGCTTCTGC  1080
Wt

Mut  1081 CCTCGGCCACCCCCGTGGCTTCCTTCCAGCTTCCCCTCTCAGGACCGAGCCCTGCCAAGT  1140
Wt

Mut  1141 CTGGCAAGAACCCGGGGAGCCCCTCTCAGGCCCAGGAGCCATCCTGA               1187
Wt
```

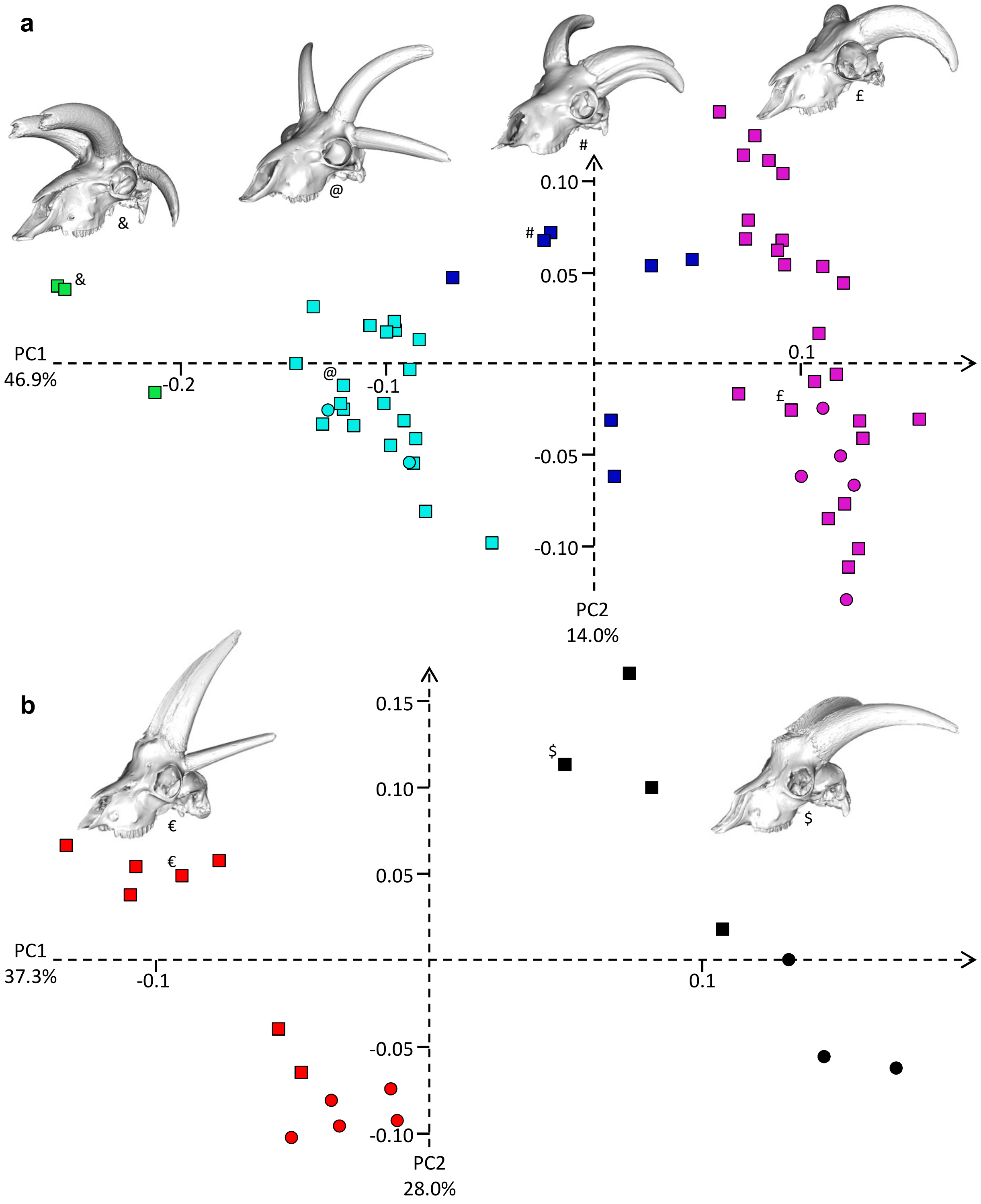


**Supplementary Figure 10. Intraspecific three-dimensional geometric morphometric analyses of 61 ovine (a) and 19 caprine (b) skulls**. Distribution of specimens along the first two axes of the PCA. The percentages of variance explained by the two main principal components are indicated on each axis. Squares and circles represent skulls from male and female animals, respectively. a) green marks: polycerate sheep with a distance between lateral horns (dlh) larger than the distance between upper horns (duh); light blue: polycerate sheep with a dlh≤duh; and blue: polycerate sheep with at least two lateral horns partially fused at their basis; purple: wild type sheep. b) red: polycerate goats; and black: wild type goats. Specimens are presented to illustrate each cluster and symbols are used to indicate their respective locations in the PCA analyses. Their respective ID in **Suppl. Table 10** are: &: Pachot_M4C_Haie; @: MNHN-ZM-AC A-12130; #: Omsr13; £: MNHN-ZM-AC A-12157; €: MNHN-ZM-AC A-12122; and $: Bresson_M2C.

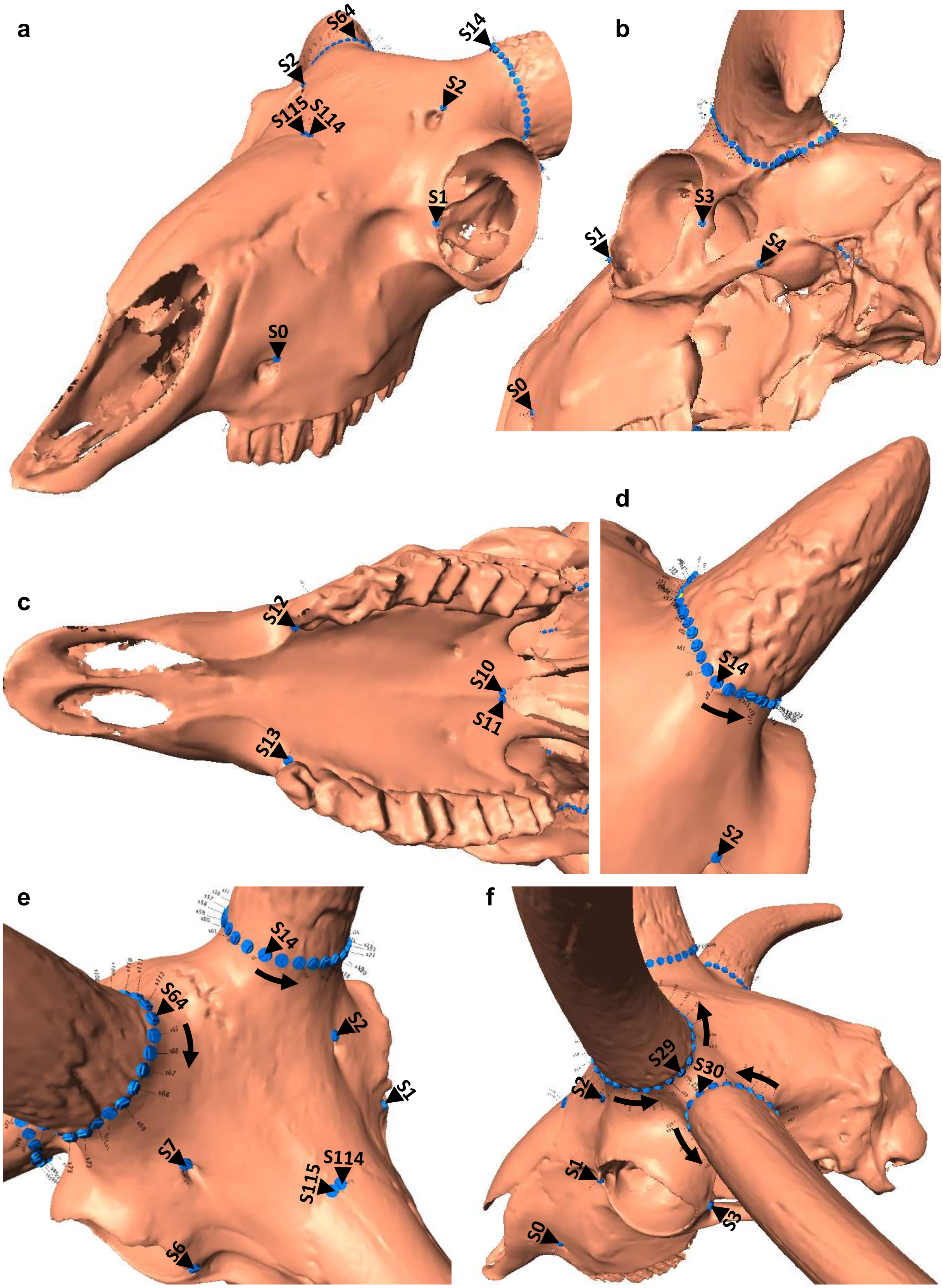


**Supplementary Figure 11. Localization of selected anatomical landmarks and sliding semi-landmarks on the skulls of wild type (a-d) and polycerate (e, f) sheep.** Arrows indicate the sense of the numbering of the sliding semi-landmarks around the neck of the cornual process. The definition of the whole landmarks is in **Suppl. Table 11.**

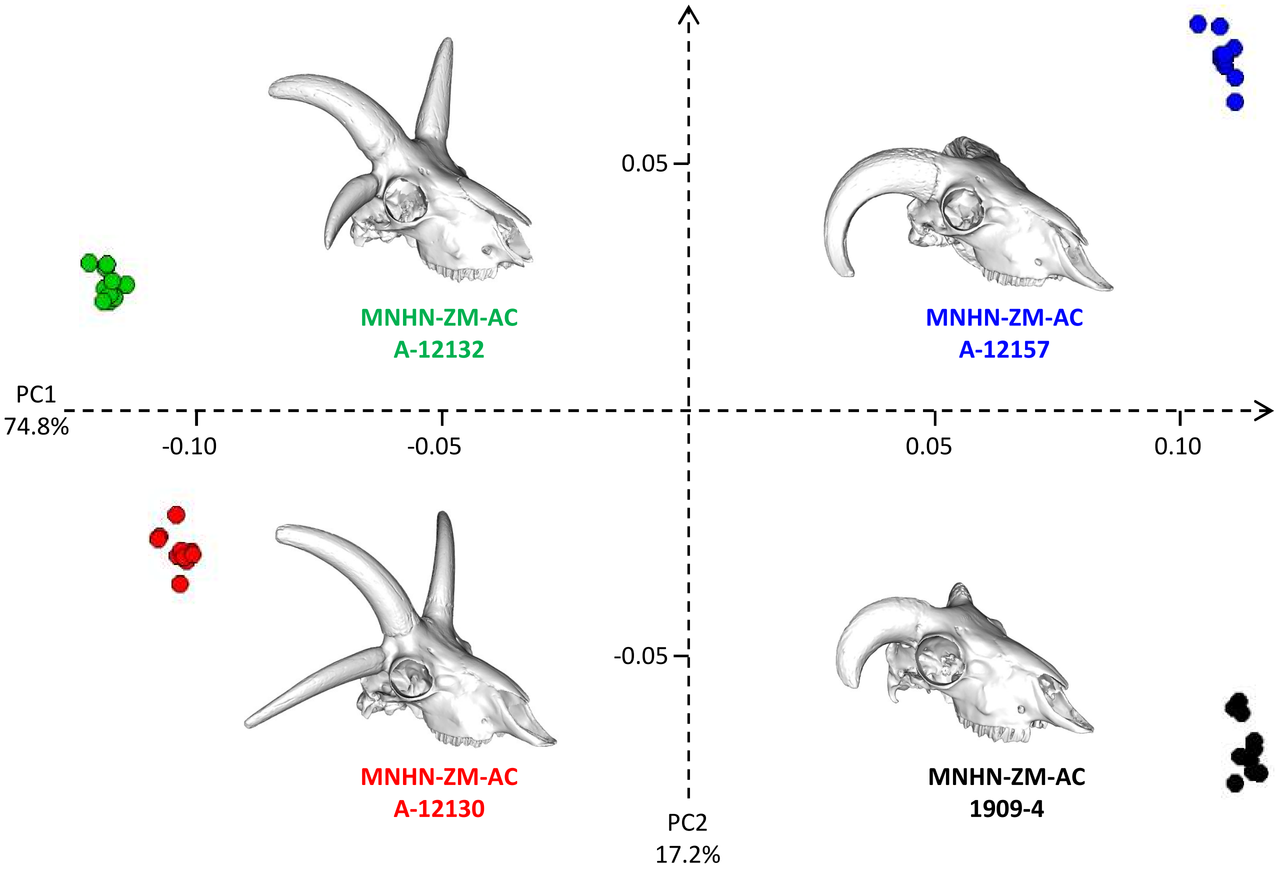


**Supplementary Figure 12. Repeatability and reproducibility of landmark placement using PCA.** A total of 116 landmarks and sliding semi-landmarks were placed ten times independently on the skulls of two polycerate and two control male sheep sampled between 1852 and 1909 in Tunisia (MNHN-ZM-AC A-12130, A12132 and 1909-4) and neighboring Algeria (MNHN-ZM-AC A12157; see **Suppl. Table 10**). The percentages of variance explained by the two main principal components are indicated on each axis. Each color corresponds to a specimen. For each skull, the inter-specimen variation on the two main axes is lower than the intra-specimen error due to differences between landmark digitalization, showing that our landmark configuration is relevant to describe shape variation within our sample.

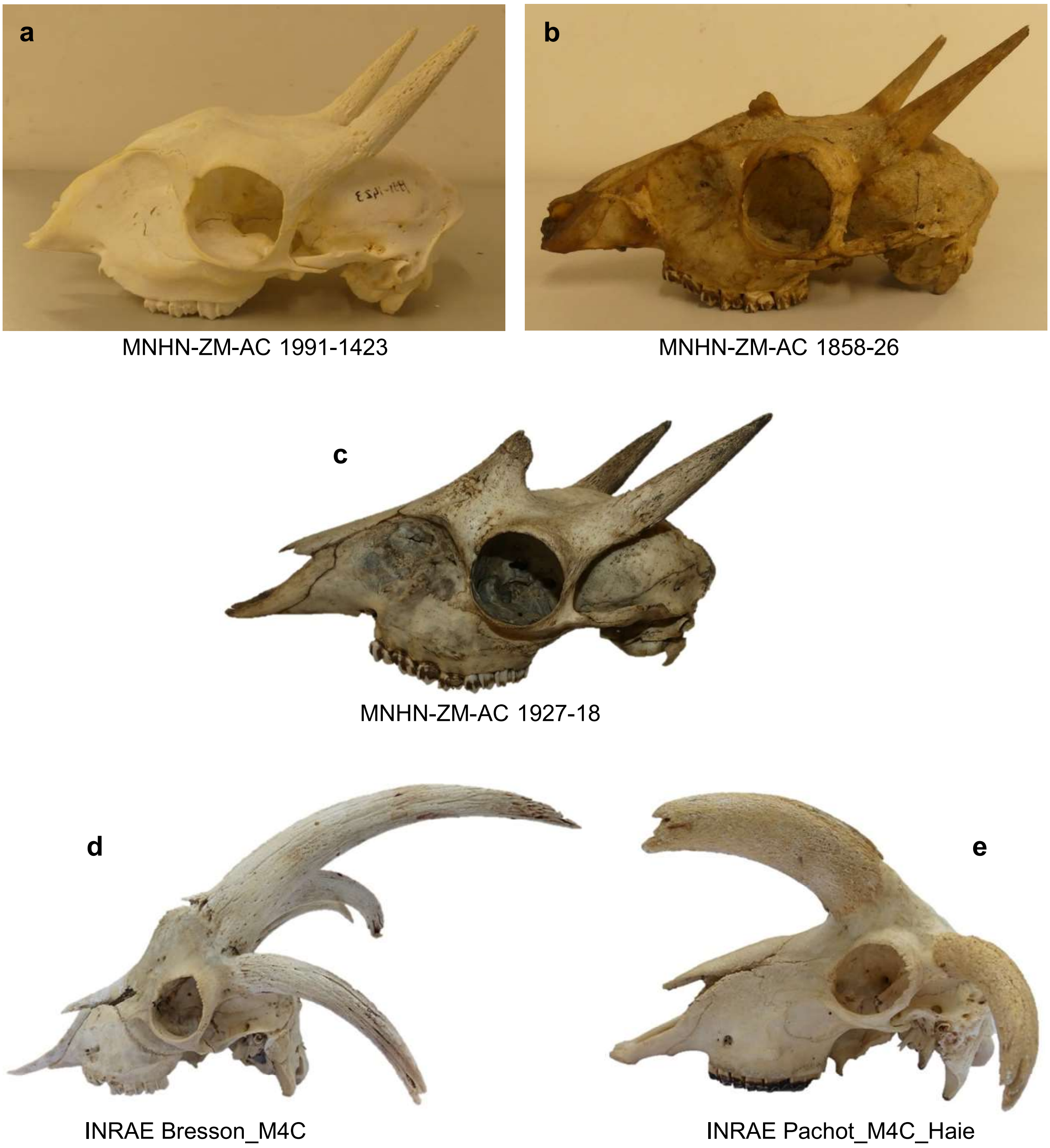


**Supplementary Figure 13. Comparison between the skulls of *Tetracerus quadricornis* and polycerate *Capra hircus* and *Ovis aries* individuals.** a-c) skulls of males *T. quadricornis* of growing age. d, e) skulls of polycerate goat (*C. hircus ;* in mirror view) and sheep (*O. aries*) males, respectively. The ID of each specimen in the MNHN and INRAE collections is indicated on the panel. Two of the three recognized subspecies of *T. quadricornis* have additional anterior horns (Groves, 2003; Leslie and Sharma, 2009), which start to develop at 10–14 months of age in captivity (Sharma *et al.* 2005). Most male *T. quadricornis* have anterior horns $^{1}/_{4}$ to $^{2}/_{3}$ the length of their posterior horns (Sharma *et al.* 2005). Note the differences in relative placement and relative size of the anterior and posterior pairs of horns in *T. quadricornis* versus polycerate *C. hircus* and *O. aries*. Specimens MNHN-ZM-AC 1991-1423, 1858-26 and 1927-18 belong to the *Collection d'anatomie comparée du Muséum National d'Histoire Naturelle*, Paris, France while specimens Bresson_M4C and Pachot_M4C_Haie originate from the collection of INRAE UMR1313 GABI, Jouy-en-Josas, France.

**Supplementary Table 1. Illumina OvineHD Beadchip genotypes used for mapping the *POLYCERATE* locus in sheep.** Wt: "wild type" refers to two normal horns or two scurs. "Polycerate" refers to the existence of more than two horns or scurs. *: In Greyvenstein *et al.* 2016, individual D-003 was mistakenly registered as polycerate while it has only two horns.

| No | Breed or population | Country of origin of the breed | Origin of the samples | Number of animals | | |
|---|---|---|---|---|---|---|
| | | | | **All** | **Wt** | **Polycerate** |
| 1 | Jacob | UK | Kijas *et al.* 2016 | 130 | 57 | 73 |
| 2 | Navajo-Churro | USA | Kijas *et al.* 2016 | 27 | 14 | 13 |
| 3 | Damara | Namibia | Greyvenstein *et al.* 2016 | 41 | 16* | 25* |
| **Total** | | | | **198** | **87** | **111** |

**Supplementary Table 2. Details on individuals considered for IBD mapping of the *POLYCERATE* locus in sheep.** Illumina OvineHD Beadchip genotyping data have been downloaded from the Dryad repository (datadryad.org/stash/dataset/doi:10.5061/dryad.1p7sf and datadryad.org/stash/dataset/doi:10.5061/dryad.6t34b) and phenotypic information have been obtained from the corresponding authors of the related articles: Kijas *et al.* 2016 and Greyvenstein *et al.* 2016. Breed : D. =Damara, J.=Jacob, and N.=Navajo-Churro; Ph. :Phenotype.

| ID | Breed | Ph. | ID | Breed | Ph. | ID | Breed | Ph. | ID | Breed | Ph. | ID | Breed | Ph. |
|---|---|---|---|---|---|---|---|---|---|---|---|---|---|---|
| D-003 | D. | 0 | H102 | J. | 0 | H36 | N. | 0 | H35 | J. | 1 | H115 | J. | 1 |
| D-008 | D. | 0 | H106 | J. | 0 | H37 | N. | 0 | H47 | J. | 1 | H116 | J. | 1 |
| D-010 | D. | 0 | H112 | J. | 0 | H42 | N. | 0 | H48 | J. | 1 | H118 | J. | 1 |
| D-011 | D. | 0 | H114 | J. | 0 | H43 | N. | 0 | H50 | J. | 1 | H121 | J. | 1 |
| D-014 | D. | 0 | H117 | J. | 0 | H45 | N. | 0 | H51 | J. | 1 | H122 | J. | 1 |
| D-018 | D. | 0 | H119 | J. | 0 | H52 | N. | 0 | H54 | J. | 1 | H123 | J. | 1 |
| D-023 | D. | 0 | H120 | J. | 0 | H58 | N. | 0 | H55 | J. | 1 | H124 | J. | 1 |
| D-076 | D. | 0 | H125 | J. | 0 | Black | D. | 1 | H57 | J. | 1 | H131 | J. | 1 |
| D-100 | D. | 0 | H126 | J. | 0 | Brown | D. | 1 | H59 | J. | 1 | H132 | J. | 1 |
| D-101 | D. | 0 | H127 | J. | 0 | D-004 | D. | 1 | H61 | J. | 1 | H133 | J. | 1 |
| D-103 | D. | 0 | H128 | J. | 0 | D-005 | D. | 1 | H62 | J. | 1 | H136 | J. | 1 |
| D-104 | D. | 0 | H129 | J. | 0 | D-013 | D. | 1 | H64 | J. | 1 | H138 | J. | 1 |
| DD11 | D. | 0 | H130 | J. | 0 | DD01 | D. | 1 | H71 | J. | 1 | H139 | J. | 1 |
| DD12 | D. | 0 | H134 | J. | 0 | DD02 | D. | 1 | H72 | J. | 1 | H143 | J. | 1 |
| DD15 | D. | 0 | H135 | J. | 0 | DD04 | D. | 1 | H76 | J. | 1 | H145 | J. | 1 |
| DD16 | D. | 0 | H137 | J. | 0 | DD05 | D. | 1 | H77 | J. | 1 | H146 | J. | 1 |
| H7 | J. | 0 | H140 | J. | 0 | DD06 | D. | 1 | H78 | J. | 1 | H147 | J. | 1 |
| H17 | J. | 0 | H141 | J. | 0 | DD07 | D. | 1 | H80 | J. | 1 | H151 | J. | 1 |
| H21 | J. | 0 | H142 | J. | 0 | DD08 | D. | 1 | H83 | J. | 1 | H155 | J. | 1 |
| H34 | J. | 0 | H144 | J. | 0 | DD09 | D. | 1 | H87 | J. | 1 | H158 | J. | 1 |
| H40 | J. | 0 | H148 | J. | 0 | DD10 | D. | 1 | H88 | J. | 1 | H159 | J. | 1 |
| H46 | J. | 0 | H149 | J. | 0 | DD13 | D. | 1 | H90 | J. | 1 | H163 | J. | 1 |
| H65 | J. | 0 | H150 | J. | 0 | DD14 | D. | 1 | H91 | J. | 1 | H164 | J. | 1 |
| H66 | J. | 0 | H152 | J. | 0 | DD17 | D. | 1 | H92 | J. | 1 | H167 | J. | 1 |
| H67 | J. | 0 | H153 | J. | 0 | DD18 | D. | 1 | H93 | J. | 1 | H170 | J. | 1 |
| H68 | J. | 0 | H154 | J. | 0 | DD20 | D. | 1 | H95 | J. | 1 | H2 | N. | 1 |
| H69 | J. | 0 | H156 | J. | 0 | DR02 | D. | 1 | H96 | J. | 1 | H8 | N. | 1 |
| H70 | J. | 0 | H157 | J. | 0 | DR03 | D. | 1 | H97 | J. | 1 | H10 | N. | 1 |
| H73 | J. | 0 | H160 | J. | 0 | DR04 | D. | 1 | H98 | J. | 1 | H11 | N. | 1 |
| H74 | J. | 0 | H161 | J. | 0 | DR05 | D. | 1 | H99 | J. | 1 | H12 | N. | 1 |
| H75 | J. | 0 | H162 | J. | 0 | DS02 | D. | 1 | H100 | J. | 1 | H14 | N. | 1 |
| H79 | J. | 0 | H166 | J. | 0 | DS03 | D. | 1 | H103 | J. | 1 | H28 | N. | 1 |
| H81 | J. | 0 | H169 | J. | 0 | H5 | J. | 1 | H104 | J. | 1 | H31 | N. | 1 |
| H82 | J. | 0 | H1 | N. | 0 | H13 | J. | 1 | H105 | J. | 1 | H38 | N. | 1 |
| H84 | J. | 0 | H4 | N. | 0 | H16 | J. | 1 | H107 | J. | 1 | H44 | N. | 1 |
| H85 | J. | 0 | H9 | N. | 0 | H20 | J. | 1 | H108 | J. | 1 | H56 | N. | 1 |
| H86 | J. | 0 | H15 | N. | 0 | H22 | J. | 1 | H109 | J. | 1 | H60 | N. | 1 |
| H89 | J. | 0 | H24 | N. | 0 | H26 | J. | 1 | H110 | J. | 1 | H63 | N. | 1 |
| H94 | J. | 0 | H29 | N. | 0 | H27 | J. | 1 | H111 | J. | 1 | | | |
| H101 | J. | 0 | H30 | N. | 0 | H32 | J. | 1 | H113 | J. | 1 | | | |

**Supplementary Table 3. Details on animals phenotyped and genotyped for validating the causative mutation of the ovine *POLYCERATE* locus.** Wt: "wild type" refers to two normal horns or two scurs. "Polycerate" refers to the existence of more than two horns or scurs. Wt: wild type.

| No | Breed or Population | Origin of the breed | Origin of the samples | Number of animals | | |
|---|---|---|---|---|---|---|
| | | | | All | Wt | Polycerate |
| 1 | Hebridean | UK | Two breeders from the Netherlands | **3** | - | 3 |
| 2 | Jacob | UK | Multiple breeders from France, Switzerland and the USA | **106** | 19 | 87 |
| 3 | Manx Loaghtan | UK | Three breeders from the UK and the Netherlands | **32** | 10 | 22 |
| 4 | Ouessant admixed with Jacob | France and UK | One breeder from Paris area, France | **6** | 4 | 2 |
| 5 | Navajo-Churro | USA | Multiple breeders from the USA | **49** | 32 | 17 |
| 6 | Ovino Quadricorna | Italy | One breeder from the Province of Frosinone, Italy | **16** | 6 | 10 |
| 7 | Local population from Tunis area | Tunisia | Zoological Park of Tunis, Tunisia | **6** | 2 | 4 |
| 8 | Damara | Namibia | Three breeders from South Africa | **18** | 12 | 6 |
| **Total for sheep (*Ovis aries*)** | | | | **236** | **85** | **151** |

**Supplementary Table 4. Association between the polycerate phenotype and chromosome 2 g.132,832,249_132,832,252del allele in 236 sheep.** Wt: wild type; Del: deletion of four nucleotides.

| Phenotype | Breed or population | Number of animals per genotype | | |
|---|---|---|---|---|
| | | Wt/Wt | Wt/Del | Del/Del |
| **Wild type** | Jacob | 19 | - | - |
| | Manx Loaghtan | 10 | - | - |
| | Ouessant admixed with Jacob | 4 | - | - |
| | Navajo-Churro | 32 | - | - |
| | Ovino Quadricorna | 6 | - | - |
| | Local population from Tunis area | 2 | - | - |
| | Damara | 12 | - | - |
| **Total for wild type animals** | | **85** | **0** | **0** |
| **Polycerate** | Hebridean | - | 2 | 1 |
| | Jacob | - | 73 | 14 |
| | Manx Loaghtan | - | 17 | 5 |
| | Ouessant admixed with Jacob | - | 2 | - |
| | Navajo-Churro | - | 16 | 1 |
| | Ovino Quadricorna | - | 9 | 1 |
| | Local population from Tunis area | - | 3 | 1 |
| | Damara | - | 5 | 1 |
| **Total for polycerate animals** | | **0** | **127** | **24** |
| **Grand total** | | **85** | **127** | **24** |

**Supplementary Table 5. Analysis of nucleotide sequence conservation at the *HOXD1* exon 1 –intron 1 junction in sarcopterygians and tetrapods.** The 40 last nucleotides of *HOXD1* exon 1, the splice donor site of exon 1 (bolded), and 20 additional nucleotides of intron 1 are presented for 103 species and genome assemblies obtained from the Ensembl (www.ensembl.org; release 98) and UCSC (http://genome.ucsc.edu/) genome browser databases.

| Species | Genome Assembly | Nucleotide sequence |
|---|---|---|
| Notamacropus eugenii | macEug2 | TTTGAATGGATGAAAGTGAAAAGAAACGCCC<br>CCAAGAAAA**GT**AAGTAAACCTTAACCTTGGG |
| Sarcophilus harrisii | sarHar1 | TTTGAATGGATGAAAGTGAAAAGAAACGCCC<br>CCAAGAAAA**GT**AAGTAAACCTTAACTTTGGG |
| Monodelphis domestica | monDom5 | TTTGAATGGATGAAAGTGAAAAGGAACGCCC<br>CCAAGAAAA**GT**AAGTAAACCTTAACCTTGGG |
| Vombatus ursinus | bare-nosed_wombat_genome_assembly | TTTGAATGGATGAAAGTGAAAAGAAACGCCC<br>CCAAGAAAAGTAAGTAAGCCTTAACCTTGGG |
| Phascolarctos cinereus | phaCin_unsw_v4.1 | TTTGAATGGATGAAAGTGAAAAGAAACGCCC<br>CCAAGAAAA**GT**AAGTAAGCCTTAACCTTGGG |
| Octodon degus | OctDeg1.0 | TTTGAGTGGATGAAAGTGAAGAGGAACACCC<br>CGAAAAAAA**GT**GAGTACATGGACCTGGGAGG |
| Chrysemys picta bellii | chrPic1 | TTCGAGTGGATGAAAGTGAAAAGAAATGCGC<br>CCAAGAAAA**GT**AAGTGTGAACCTCGGCGAGG |
| Gopherus agassizii | ASM289641v1 | TTCGAGTGGATGAAAGTGAAAAGAAATGCGC<br>CCAAGAAAA**GT**AAGTGTGAACCTCGGCGAGG |
| Chelonoidis abingdonii | ASM359739v1 | TTCGAGTGGATGAAAGTGAAAAGAAATGCGC<br>CCAAGAAAA**GT**AAGTGTGAACCTCGGCAAGG |
| Canis lupus familiaris | canFam3 | TTCGAGTGGATGAAAGTGAGGAGGAGCGCCC<br>CTCGGAAAA**GT**AAGTGCGGGCCGCGGGCGGG |
| Salvator merianae | HLtupMer3 | TTCGAGTGGATGAAAGTCAAGAGGAACGCAC<br>CTCCGAAAG**GT**AAGCAGGGCAGCCTCCGGAG |
| Alligator mississippiensis | allMis1 | TTCGAGTGGATGAAGGTGAAGAGAAACGCGC<br>CCAGGAAAA**GT**AAGTCTCCAGCCGGGGGAGG |
| Crocodylus porosus | CroPor_comp1 | TTCGACTGGATGAAGGTGAAGAGAAACGCGC<br>CCAGGAAAA**GT**AAGTCTCCAGCCGGGGGGGT |
| Coturnix japonica | Coturnix_japonica_2.0 | TTCGAGTGGATGCGGATGAAGCGGAGCACGC<br>CAGGCAGAA**GT**GAGTGGGGCGGCGGGGCGCG |
| Numida meleagris | NumMel1.0 | TTCGAGTGGATGCGGATGAAGCGGAGCACGC<br>CCGGTAGAA**GT**GAGTGGGGCGGCGGGGCGCG |
| Anser brachyrhynchus | ASM259213v1 | TTCGAGTGGATGCGAGTGAAGCGGAGCCCGC<br>CCGGGAGAA**GT**GAGTGGGGCCGCGGGGCGCG |
| Pelodiscus sinensis | PelSin_1.0 | TTCGAGTGGATGAAAGTGAAGCGAAACGCAC<br>CCACGAAAA**GT**AAGTGTAACCCTCGGGGGGC |
| Meriones unguiculatus | MunDraft-v1.0 | TTCGAGTGGATGAAAGTGAAGAGGAACGCCC<br>CCAGGAAAA**GT**AAGGAGGTGGGCGCCGGGGG |
| Dipodomys ordii | dipOrd1 | TTCGAATGGATGAAAGTGAAGAGGAACGCCC<br>CTAAGAAGA**GT**AAGTACTGGGGCGTGGGGTG |
| Ornithorhynchus anatinus | ornAna2 | TTCGACTGGATGAAAGTGAAAAGAAACGCGC<br>CCAGGAAAA**GT**AAGTAGTCCTCGGGCGACCT |
| Fukomys damarensis | DMR_v1.0 | TTCGAGTGGATGAAAGTAAAGAGGAACGCCC<br>CTAAGAAAA**GT**AAGTACTTGGGCGAGGGGTG |
| Microcebus murinus | micMur2 | TTTGAGTGGATGAAAGTGAAGAGGAACGCCC<br>CTAAGAAAA**GT**AAGTACGTGGGCCTCGGGCC |
| Manis pentadactyla | manPen1 | TTTGAGTGGATGAAAGTGAAGAGGAACACCC<br>CTAAGAGAA**GT**AAGTACTTGGGCCCTGGACC |
| Otolemur garnettii | otoGar3 | TTTGAATGGATGAAAGTGAAGAGGAATGCCC<br>CTAAGAAAA**GT**AAGTTGTGGGCCTTGGATGG |
| Propithecus coquereli | Pcoq_1.0 | TTTGAGTGGATGAAAGTGAAGAGGAACGCTC<br>CTAAGAAAA**GT**AAGTAGTGGGCCTTGGACCG |

**Supplementary Table 5, continuing**

| **Species** | **Genome Assembly** | **Nucleotide sequence** |
|---|---|---|
| Prolemur simus | Prosim_1.0 | TTTGAGTGGATGAAAGTGAAGAGGAACGCCC CTAAGAAAA**GT**AAGTCCGTGGCCTTGGTCG |
| Mus spretus | SPRET_EiJ_v1 | TTCGAGTGGATGAAAGTGAAGAGGAACGCCC CCAAGAAAA**GT**AAGGAGGTGGGCGCTGAGGG |
| Mus caroli | CAROLI_EIJ_v1.1 | TTCGAGTGGATGAAAGTGAAGAGGAACGCCC CCAAGAAAA**GT**AAGGAGGTGGGCGCTGAGGG |
| Mus musculus | GRCm38.p6 | TTCGAGTGGATGAAAGTGAAGAGGAACGCCC CCAAGAAAA**GT**AAGGAGGTGGGCGCTGAGGG |
| Mus pahari | PAHARI_EIJ_v1.1 | TTCGAGTGGATGAAAGTGAAGAGGAACGCCC CCAAGAAAA**GT**AAGGAGGTGGGCGCTGTGGG |
| Rattus norvegicus | rn6 | TTCGAGTGGATGAAAGTGAAGAGGAACGCCC CCAAGAAAA**GT**AAGGAGGTGGGCGCTGGGGG |
| Cricetulus griseus | criGriChoV2 | TTCGAGTGGATGAAAGTGAAGAGGAACGCCC CCAGGAAAA**GT**AAGGAGGTGGGCGCTGGAGG |
| Peromyscus maniculatus bairdii | HU_Pman_2.1 | TTCGAGTGGATGAAAGTCAAGAGGAACGCCC CCAAGAAAA**GT**AAGGAGGTGGGCGCTGGAGG |
| Mesocricetus auratus | MesAur1.0 | TTCGAGTGGATGAAAGTGAAGAGGAACGCCC CCAAGAAAA**GT**AAGAAGGTGGGCGCTGGAGG |
| Microtus ochrogaster | MicOch1.0 | TTCGAGTGGATGAAAGTGAAGAGGAACGCCC CCAAGAAAA**GT**AAGGAGACGGACGCTGGAGG |
| Mus spicilegus | MUSP714 | TTCGAGTGGATGAAAGTGAAGAGGAACGCCC CCAAGAAAA**GT**AAGGGGGGGGGCGGGGGGG |
| Cavia porcellus | cavPor3 | TTTGAGTGGATGAAAGTAAAGAGGAACGCCC CTAAGAAAA**GT**AAGTACGTGGGCGCTGGGTG |
| Chinchilla lanigera | ChiLan1.0 | TTCGAGTGGATGAAAGTAAAGAGGAACGCCT CTAAGAAAA**GT**AAGTACAGGGCGCTGGTGCA |
| Erinaceus europaeus | eriEur2 | TTCGAGTGGATGAAAGTGAAGCGCAGCGCCC CGAAGAGAA**GT**AAGTCGCCGGCCCCCGGGGC |
| Nannospalax galili | S.galili_v1.0 | TTCGAGTGGATGAAAGTGAAGAGGAACGCCC AGAAGAAAA**GT**AAGTGGGTGGGCGCTGAGGG |
| Ictidomys tridecemlineatus | speTri2 | TTCGAGTGGATGAAAGTGAAGAGGAACGCCC CTAAGAAAA**GT**AAGTCGCTGGGCTCTGGATG |
| Urocitellus parryii | ASM342692v1 | TTCGAGTGGATGAAAGTGAAGAGGAACGCCC CTAAGAAAA**GT**AAGTCGCTGGGCTCTGGATG |
| Spermophilus dauricus | ASM240643v1 | TTCGAGTGGATGAAAGTGAAGAGGAACGCCC CTAAGAAAA**GT**AAGTCGCTGGGCGCTGGATG |
| Heterocephalus glaber | hetGla2 | TTCGAGTGGATGAAAGTGAAGAGGAATGCCC CTAAGAAAA**GT**AAGTACTTGGGTGCCGGGTG |
| Tupaia belangeri | tupBel1 | TTTGAGTGGATGAAAGTGAAGAGGAATGCCC CTAAGAAAA**GT**AAGTCCTTGGCTTCTGATGC |
| Ochotona princeps | ochPri3 | TTCGAGTGGATGAAAGTGAAAAGGAACGCTC CTAAGAAAA**GT**AAGTGAAGTAGGGTTTAGAC |
| Myotis lucifugus | myoLuc2 | TTCGAGTGGATGAAAGTGAAGAGGAACGCCC CCAAGAAAA**GT**GAGTACTCGAGTCTTGGACG |
| Oryctolagus cuniculus | oryCun2 | TTCGAGTGGATGAAAGTGAAAAGAAACGCCC CAAAGAAAA**GT**AAGTACTGAGCTTTGGATGC |
| Sus scrofa | susScr11 | TTCGAGTGGATGAAAGTGAAGAGGAACGCCC CGAAGAAAA**GT**AAGTACTTGGGCTCTCGACG |
| Vicugna pacos | vicPac2 | TTCGAGTGGATGAAAGTGAAAAGGAACGCCC CGAAGAAAA**GT**AAGTACTTGGGCCCTGGACC |
| Tursiops truncatus | turTru2 | TTCGAGTGGATGAAAGTGAAGAGGAACGCCC CGAAAAAAA**GT**AAGTGCTTGGGCCTTGGACG |
| Ovis aries | oviAri4 | TTCGAGTGGATGAAAGTGAAGAGGAACGCGC CGAAGAAAA**GT**AAGTACTTGAGCCTTGGACG |
| Capra hircus | ARS1 | TTCGAGTGGATGAAAGTGAAGAGGAACGCGC CGAAGAAAA**GT**AAGTACTTGAGCCTTGGACG |

**Supplementary Table 5, continuing**

| Species | Genome Assembly | Nucleotide sequence |
|---|---|---|
| Bos taurus | bosTau9 | TTCGAGTGGATGAAAGTGAAGAGGAACGCCC CGAAGAAAA**GT**AAGTACTTGGGCCTTGGACG |
| Bison bison bison | Bison_UMD1.0 | TTCGAGTGGATGAAAGTGAAGAGGAACGCCC CAAAGAAAA**GT**AAGTACTTGGGCCTTGGACG |
| Balaenoptera acutorostrata scammoni | balAcu1 | TTCGAGTGGATGAAAGTGAAGAGGAACGCCC CGAAGAAAA**GT**AAGTGCTTGGGCCTTGGACG |
| Felis catus | felCat9 | TTCGAGTGGATGAAAGTGAAGAGGAACGCCC CTAAGAAAA**GT**AAGTATCTGGGCCTTGGACG |
| Panthera pardus | PanPar1.0 | TTCGAGTGGATGAAAGTGAAGAGGAACGCCC CTAAGAAAA**GT**AAGTATCTGGGCCTTGGACG |
| Pteropus vampyrus | pteVam1 | TTCGAGTGGATGAAAGTGAAGAGGAACGCCC CGAAGAAAA**GT**AAGTATTTGAGCCATGGACG |
| Mustela putorius furo | musFur1 | TTCGAGTGGATGAAAGTGAAGAGGAACGCCC CTAGGAAAA**GT**AAGTGCTTGGGCCTCGGACC |
| Neovison vison | NNQGG.v01 | TTTGAGTGGATGAAAGTGAAGAGGAACGCCC CTAGGAAAA**GT**AAGTGCTTGGGCCTCGGACC |
| Ailuropoda melanoleuca | ailMel1 | TTCGAGTGGATGAAAGTGAAGAGGAACGCCC CTAGGAAAA**GT**AAGTACTTGGGCCTCGGACC |
| Ursus americanus | ASM334442v1 | TTCGAGTGGATGAAAGTGAAGAGGAACGCCC CTAGGAAAA**GT**AAGTACTTGGGCCTCGGACC |
| Loxodonta africana | loxAfr3 | TTCGAGTGGATGAAAGTGAAGAGGAACGCCC CTAAGAAAA**GT**AAGTACTTGGGTCTTGGACG |
| Trichechus manatus latirostris | triMan1 | TTCGAGTGGATGAAAGTGAAGAGGAACGCCC CTAAGAAAA**GT**AAGTACTTGGGCCTTGGACG |
| Procavia capensis | proCap1 | TTCGAGTGGATGAAAGTGAAGAGGAACGCGC CTAAGAAAA**GT**AAGTATTTGGGCCTTGGACA |
| Dasypus novemcinctus | dasNov3 | TTCGAGTGGATGAAAGTGAAGAGGAGCGCCT CTAGGAAAA**GT**AAGTGTTGGGCCTTAGACAG |
| Equus caballus | equCab3 | TTCGAGTGGATGAAAGTGAAGAGGAACGCCC CCAAGAAAA**GT**GAGTACTTGGGCCGTGGATG |
| Equus asinus asinus | ASM303372v1 | TTCGAGTGGATGAAAGTGAAGAGGAACGCCC CCAAGAAAA**GT**GAGTACTTGGGCCGTGGATG |
| Galeopterus variegatus | galVar1 | TTCGAGTGGATGAAAGTGAAGAGGAACGCCC CTAAGAAAA**GT**AAGTACTTGGGCCGTGGATG |
| Castor canadensis | C.can_genome_v1.0 | TTCGAGTGGATGAAAGTGAAGAGGAACGCCC CTAAGAAAA**GT**AAGTACGCAGGCGTTGGATG |
| Callithrix jacchus | calJac3 | TTCGAGTGGATGAAAGTGAAGAGGAACGCCC CTAAGAAAA**GT**AAGTCCTCGGGCCTTGGATG |
| Cebus capucinus imitator | Cebus_imitator-1.0 | TTCGAGTGGATGAAAGTGAAGAGGAACGCCC CTAAGAAAA**GT**AAGTCCGCGGGCCTTGGATG |
| Saimiri boliviensis boliviensis | saiBol1 | TTCGAGTGGATGAAAGTGAAGAGGAACGCTC CTAAGAAAA**GT**AAGTCCGCGGGCCTTGGATG |
| Aotus nancymaae | Anan_2.0 | TTCGAGTGGATGAAAGTGAAGAGGAACGCCC CTAAGAAAA**GT**AAGTCCGCGGGCCTTGGATG |
| Homo sapiens | hg38 | TTCGAGTGGATGAAAGTGAAGAGGAATGCCT CTAAGAAAG**GT**AAGTCCGCGGGCCTTGGATG |
| Gorilla gorilla gorilla | gorGor5 | TTCGAGTGGATGAAAGTGAAGAGGAATGCCT CTAAGAAAG**GT**AAGTCCGCGGGCCTTGGATG |
| Pongo abelii | ponAbe3 | TTCGAGTGGATGAAAGTGAAGAGGAATGCCT CTAAGAAAG**GT**AAGTCCGCGGGCCTTGGATG |
| Pan troglodytes | panTro6 | TTCGAGTGGATGAAAGTGAAGAGGAATGCCT CTAAGAAAG**GT**AAGTCCGCGGGCCTTGGATA |
| Pan paniscus | panPan2 | TTCGAGTGGATGAAAGTGAAGAGGAATGCCT CTAAGAAAG**GT**AAGTCCGCGGGCCTTGGATA |
| Nomascus leucogenys | nomLeu3 | TTCGAGTGGATGAAAGTGAAGAGGAATGCCT CTAAGAAAG**GT**AAGTCCGCGGTCCTTGGATG |

**Supplementary Table 5, continuing**

| Species | Genome Assembly | Nucleotide sequence |
|---|---|---|
| Cercocebus atys | Caty_1.0 | TTCGAGTGGATGAAAGTGAAGAGGAACGCCT<br>CTAAGAAAA**GT**AAGTCCGCGGGCCTTGGATG |
| Piliocolobus tephrosceles | ASM277652v2 | TTCGAGTGGATGAAAGTGAAGAGGAACGCCT<br>CTAAGAAAA**GT**AAGTCCGCGGGCCTTGGATG |
| Theropithecus gelada | Tgel_1.0 | TTCGAGTGGATGAAAGTGAAGAGGAACGCCT<br>CTAAGAAAA**GT**AAGTCCGCGGGCCTTGGATG |
| Mandrillus leucophaeus | Mleu.le_1.0 | TTCGAGTGGATGAAAGTGAAGAGGAACGCCT<br>CTAAGAAAA**GT**AAGTCCGCGGGCCTTGGATG |
| Rhinopithecus bieti | ASM169854v1 | TTCGAGTGGATGAAAGTGAAGAGGAACGCCT<br>CTAAGAAAA**GT**AAGTCCGCGGGCCTTGGATG |
| Colobus angolensis palliatus | Cang.pa_1.0 | TTCGAGTGGATGAAAGTGAAGAGGAACGCCT<br>CTAAGAAAA**GT**AAGTCCGCGGGCCTTGGATG |
| Rhinopithecus roxellana | rhiRox1 | TTCGAGTGGATGAAAGTGAAGAGGAACGCCT<br>CTAAGAAAA**GT**AAGTCCGCGGGCCTTGGATG |
| Nasalis larvatus | nasLar1 | TTCGAGTGGATGAAAGTGAAGAGGAACGCCT<br>CTAAGAAAA**GT**AAGTCCGCGGGCCTTGGATG |
| Chlorocebus sabaeus | chlSab2 | TTCGAGTGGATGAAAGTGAAGAGGAACGCCT<br>CTAAGAAAA**GT**AAGTCCGCGGGCCTTGGATG |
| Macaca fascicularis | Macaca_fascicularis_5.0 | TTCGAGTGGATGAAAGTGAAGAGGAACGCCT<br>CTAAGAAAA**GT**AAGTCCGCGGGCTTGGATGG |
| Macaca mulatta | Mmul_10 | TTCGAGTGGATGAAAGTGAAGAGGAACGCCT<br>CTAAGAAAA**GT**AAGTCCGCGGGCTTGGATGG |
| Macaca nemestrina | Mnem_1.0 | TTCGAGTGGATGAAAGTGAAGAGGAACGCCT<br>CTAAGAAAA**GT**AAGTCCGCGGGCTTGGATGG |
| Papio anubis | papAnu4 | TTCGAGTGGATGAAAGTGAAGAGGAACGCCT<br>CTAAGAAAA**GT**AAGTCCGCAGGCCTTGGATG |
| Papio hamadryas | papHam1 | TTCGAGTGGATGAAAGTGAAGAGGAACGCCT<br>CTAAGAAAA**GT**AAGTCCGCAGGCCTTGGATG |
| Sorex araneus | sorAra2 | TTCGAGTGGATGAAAGTGAAGAGGAACGCTC<br>CGAAGAAAA**GT**AAGTACTTGGCCCGTGGACT |
| Jaculus jaculus | JacJac1.0 | TTCGAGTGGATGAAAGTGAAGAGGAACGCCC<br>CGAGGAAAA**GT**GAGTGGTGGGCGATGGATGG |
| Thamnophis sirtalis | thaSir1 | TTCGACTGGATGAAAGTCAAGAGGAACGCAC<br>CCCCTAAAA**GT**AAGTGCCGGGGCTTGGTCAA |
| Notechis scutatus | TS10Xv2-PRI | TTCGAGTGGATGAAAGTCAAGAGGAACGCGC<br>TCCAGAAAA**GT**AAGTGCGGGGCTTCGTCGGC |
| Xenopus tropicalis | xenTro9 | TTTGATTGGATGAAAGTTAAAAGGAACCCAC<br>CTAAGAAAA**GT**AAGTCCTGCCAAGCTTGATA |
| Xenopus laevis | xenLae2 | TTTGATTGGATGAAAGTTAAAAGGAACCCTC<br>CTAAGAAAA**GT**AAGTTTTTATTAGCGCGAAA |
| Latimeria chalumnae | LatCha1 | TTCGATTGGATGAAGGTTAAAAGAAATCCTC<br>CCAAAACAT**GT**AAGTCTCGGTACAACAATAA |
| Nanorana parkeri | nanPar1 | TTTGATTGGATGAAAGTTAAACGGAACGCTC<br>CTAAGAAAA**GT**AAGTACAAAGAGATCCCATT |
| **Conservation** |  | ** ** ****** * * * * *<br>* ** ** |

**Supplementary Table 6. Illumina GoatSNP50 BeadChip genotypes generated for mapping the *POLYCERATE* locus in goat.** Wt : "wild type" refers to the presence of two normal horns while "polycerate" refers to the existence of more than two horns. a: in these admixed populations the polycerate phenotype originates from local ancestry.

| No | Breed or population | Origin of the breed | Origin of the samples | Number of animals | | |
|---|---|---|---|---|---|---|
| | | | | All | Wt | Polycerate |
| 1 | Local population of Massif Central admixed with African dwarf goat[a] | France | One breeder from the St Etienne area, France | 1 | - | 1 |
| 2 | Local population of Normandy admixed with Nubian goat[a] | France | Parc Animalier du Beauquet Marais, Normandy, France | 5 | 2 | 3 |
| 3 | Provençale | France | One breeder from Moustiers-Sainte-Marie, France | 44 | 31 | 14 |
| 4 | Provençale crossbred with Alpine and Saanen | France | UCEA Bressonvilliers, INRAE experimental farm, France | 16 | 11 | 5 |
| 5 | Rove admixed with Provençale | France | One breeder from Moustiers-Sainte-Marie, France | 3 | - | 3 |
| 6 | Population from the German Alps admixed with other breeds[a] | Germany | Tierpark Hamm, North Rhine-Westphalia and one breeder from Asendorf, Lower Saxony, Germany | 5 | 2 | 3 |
| 7 | Local population | Italy | One breeder from the Osimo locality, Italy | 8 | 4 | 4 |
| 8 | Local population | Macedonia | Two breeders from Radovis and Strumica municipalities, Macedonia | 3 | 1 | 2 |
| **Total** | | | | **86** | **51** | **35** |

**Supplementary Table 7. Informatin on candidate variants for polyceraty in goat and primers used for genotyping.** Genotyping was done by PCR and Sanger sequencing on a panel of 5 case-control pairs for all the variants except g.115,652,290_116,155,699delins137kb. For the latter, the genotyping was done on a panel of 77 case and 355 control animals by PCR and electrophoresis, using two primers specific of the wild type (Wt) and mutant (Mut) alleles, respectively and one primer in common. The expected size of the wild type and mutant amplicons are 226 and 197 bp respectively (see Supplementary Fig. 4). Positions refer to ARS1 goat genome assembly. Conservation : "Yes" indicates that the variant affects at least one nucleotide located within a constrained element among 103 eutherian mammals genomes according to Ensembl (www.ensembl.org; release 98) EPO-Low-Coverage track and that this nucleotide is entirely conserved among the 103 species. Genotyping : "Eliminated" refers to variants for which the derived allele was not perfectly associated with polyceraty in the panel studied. *) Despite five trials with distinct primer pairs we failed to genotype these variants because of lack of PCR amplification or amplification of multiple segments. Therefore these variants are considered as retained by default. NC: not communicated. Cons. : conservation.

| Variant | Primer sequences | Cons. | Genotyping |
|---|---|---|---|
| g.115,143,545G>A | CCTTGGCTTGCCTGAATCTC<br>CCAGCCGTATCACAGTCAGA | No | Eliminated |
| g.115,166,967_115,166,968insT | CGGGAGCAAAAGCCTACATG<br>CTGGAGATGGACGGTCATGA | No | Eliminated |
| g.115,186,815G>A | CCCGGTTTGATTCCTAGGTT<br>CTTGCAAGATCCTCCGTTTC | No | Eliminated |
| g.115,213,759G>C | CCTGGCTTGGAGAATTTTGA<br>TGAAGGAACAAGGCTGACCT | No | **Retained** |
| g.115,221,938C>T | ATCACAGCGCTATGAGGGTT<br>ACCCTTATTGTGAAATAGGCAGA | No | **Retained** |
| g.115,223,871_115,223,871insA | ACTCCAGTAAGTGCAGTCTTCA<br>AGAAAGGAGGCAGCTGTTCT | No | Eliminated |
| g.115,232,184T>C | CCTGCAGTGTTATTTGGGCA<br>GGCATACATGTGTCAGGCTG | No | **Retained** |
| g.115,269,955A>G | TCAGTGCTCAGCCTTCTTCA<br>GTGCCACAACAGCTATCTTCT | No | **Retained** |
| g.115,283,205G>A | GCCTGTTCAGAGAATGCCTG<br>TTCTCCCTTCTGGCTCTGTG | No | Eliminated |
| g.115,285,788C>T | AAGGGCTGCTGTGTCACTAT<br>AATCAACAGTCTCGGCTCCA | No | Eliminated |
| g.115,319,352A>G | TGAACAAATCCATCAGATGAGTC<br>TCCCTTTCTTTTGTTTCAGCA | No | Eliminated |
| g.115,337,441C>T | TCGTGTCCAGTCCTAAGCAT<br>GCCATTGACGGATTGAGGAA | No | **Retained** |
| g.115,343,411del* | NC | No | **Retained by default** |
| g.115,349,841G>T | TTACAAGCCCAACAGTAACTC<br>ATGGGCCGTGTGGGATCA | No | Eliminated |
| g.115,378,261G>A | TCCTTGTTTCATAATGAGTACT<br>ATAATTTGACCAAAGTCAATGCA | No | **Retained** |
| g.115,393,837A>G | GTCTCAGTTCAAAATGGCGGA<br>GTCCCTTTGTCAGATCCCCA | No | **Retained** |
| g.115,401,255C>A | ACTTGGAATGCAGAGACAGC<br>TTCCCGTTTGATCTTTCCGC | No | **Retained** |
| g.115,406,825A>T | CGCTAACTTCGAAACACTTTGAC<br>CCCAGCGTCCTAAGTGAACT | No | Eliminated |
| g.115,409,283A>G | GTCCACAACCAGTCAACACC<br>AATCTTCCCTCCCCAGTCAC | No | **Retained** |
| g.115,417,698C>A | CTGGGGAAGGTGTTATGGGT<br>TGGGAATCTTTGAGGGCCAT | No | Eliminated |

**Supplementary Table 7, continued**

| Variant | Primer sequences | Cons. | Genotyping |
|---|---|---|---|
| g.115,422,038T>G | GGAGTCTAGAACCCTGAACCC<br>TCTCCAGCCGTTATCCCTTC | No | Eliminated |
| g.115,459,737A>G | GGCCTGAGATCCTGTGGTAA<br>CCCTGCGCTATTCAGTAGGA | No | Eliminated |
| g.115,462,622del | GGCAGCTATAGAATACACTGTGC<br>TGCCATTCTTGTAGGTCCGT | No | Eliminated |
| g.115,465,345T>A | TAGCACCATGACACCACACT<br>CCTCCATACACAACACAGCG | No | Eliminated |
| g.115,474,151A>C | CCATTGCTTTTCTGTGCCCT<br>GGGAAAGCTTGGACACGTTT | No | Eliminated |
| g.115,560,482T>A* | NC | No | **Retained by default** |
| g.115,566,277G>A | ATCAGTGTGACAATTGCCCG<br>AGATTTCCAGCACACTCCGA | No | **Retained** |
| g.115,570,129_115,570,130 delinsCG | TCTTTGCTTTCCCCTTCCCA<br>CCAGTCAGTACCCACCACAT | No | Eliminated |
| g.115,620,572A>C | GTTGTATGCGCTGACCTGAG<br>TCCTAGCCAAGTGATCCTGG | No | Eliminated |
| g.115,621,441C>T | AAGGGCAGTGGACAACATCA<br>AACTCCAAGGTCCTCTGTGC | No | Eliminated |
| g.115,629,515G>T | CTGGGTCCTTGGGCTCTTTA<br>GTCGCCGTGAGATTCAGTTC | No | **Retained** |
| g.115,639,463G>C* | NC | No | **Retained by default** |
| g.115,643,117C>T | TTCTGTCCCCTGCATGTCTT<br>GGCTTACATTGTCCCACTGG | No | Eliminated |
| g.115,644,117T>A* | NC | No | **Retained by default** |
| g.115,646,033T>C | GAACAACAAGAACGTGGGCT<br>TTGGCTGGAGAGTTGGATTT | No | Eliminated |
| g.115,652,290_116,155,699 delins137kb | CTTTCAAAGCAGTGTAATAGGA(Wt)<br>GCATGTCTACATACTTATTTAAG(Mut)<br>AGATATAACAAGAGCAAGACTG(Common) | Yes | **Retained** |

**Supplementary Table 8. Details on wild type and polycerate goats phenotyped and genotyped for candidate mutations.** Wt: "wild type" refers to the presence of two normal horns while "polycerate" refers to the existence of more than two horns. a: In these admixed populations the polycerate phenotype originates from local ancestry. The whole panel was genotyped for mutation g.115,652,290_116,155,699delins137kb while numbers between brackets refer to individuals genotyped for the other candidate mutations detailed in **Suppl. table 7.**

| No | Breed or population | Origin of the breed | Origin of the samples | Number of animals | | |
|---|---|---|---|---|---|---|
| | | | | All | Wt | Polycerate |
| 1 | Alpine | France | Multiple breeders from France | 30 | 30 | - |
| 2 | Corse | France | Multiple breeders from France | 30 | 30 | - |
| 3 | Local population of Massif Central admixed with African dwarf goat[a] | France | One breeder from the St Etienne area, France | 5 | - | 5 |
| 4 | Local population of Normandy admixed with Nubian goat[a] | France | Parc Animalier du Beauquet Marais, Normandy, France | 7 (2) | 2 (1) | 5 (1) |
| 5 | Poitevine | France | Multiple breeders from France | 30 | 30 | - |
| 6 | Provençale | France | One breeder from Moustiers-Sainte-Marie, France | 76 (2) | 45 (1) | 31 (1) |
| 7 | Provençale admixed with Alpine and Saanen | France | UCEA Bressonvilliers, INRAE experimental farm, France | 16 | 11 | 5 |
| 8 | Pyrénéenne | France | Multiple breeders from France | 30 | 30 | - |
| 9 | Rove admixed with Provençale | France | One breeder from Moustiers-Sainte-Marie, France | 4 | - | 4 |
| 10 | Population from the German Alps admixed with other breeds[a] | Germany | Tierpark Hamm, North Rhine-Westphalia and one breeder from Asendorf, Lower Saxony, Germany | 20 (2) | 9 (1) | 11 (1) |
| 11 | Appenzell | Switzerland | Multiple breeders from the canton of Appenzell, Switzerland | 5 | 5 | - |
| 12 | Booted goat | Switzerland | Multiple breeders from the canton of St. Gallen, Switzerland | 10 | 10 | - |
| 13 | Capra grigia | Switzerland | Multiple breeders from the canton of Ticino, Switzerland | 10 | 10 | - |
| 14 | Grisons striped | Switzerland | Multiple breeders from the canton of Grisons, Switzerland | 10 | 10 | - |
| 15 | Nera verzasca | Switzerland | Multiple breeders from the canton of Ticino, Switzerland | 10 | 10 | - |
| 16 | Peacock goat | Switzerland | Multiple breeders from the canton of St. Gallen, Switzerland | 10 | 10 | - |
| 17 | Saanen | Switzerland | Multiple breeders from France | 30 | 30 | - |

**Supplementary Table 8, continued**

| No | Breed or population | Origin of the breed | Origin of the samples | Number of animals | | |
|---|---|---|---|---|---|---|
| | | | | **All** | **Wt** | **Polycerate** |
| 18 | Toggenburg | Switzerland | Multiple breeders from the canton of St. Gallen, Switzerland | 10 | 10 | - |
| 19 | Valais blackneck | Switzerland | Multiple breeders from the canton of Valais, Switzerland | 10 | 10 | - |
| 20 | Local population | Italy | One breeder from the Osimo locality, Italy | 23 (2) | 10 (1) | 13 (1) |
| 21 | Local population | Croatia | Four breeders from the Benkovac municipality, Croatia | 22 | 22 | - |
| 22 | Local population | Macedonia | Two breeders from Radovis and Strumica municipalities, Macedonia | 7 (2) | 4 (1) | 3 (1) |
| 23 | Angora | Turkey | Multiple breeders from France and Germany | 17 | 17 | - |
| 24 | Boer | South Africa | Multiple breeders from Switzerland | 10 | 10 | - |
| **Total** | | | | **432 (10)** | **355 (5)** | **77 (5)** |

**Supplementary Table 9. Association between the polycerate phenotype and chromosome 2 g.115,652,290_116,155,699delins137kb in 432 goats.** Wt: wild type; Indel: deletion of 503 kb and insertion of 137 kb. a: In these admixed populations the polycerate phenotype originates from local ancestry.

| Phenotype | Breed/Population | Number of animals per genotype | | |
|---|---|---|---|---|
| | | Wt/Wt | Wt/Indel | Indel/Indel |
| Wild type | Alpine | 30 | - | - |
| | Corse | 30 | - | - |
| | Local population of Normandy admixed with Nubian goat[a] | 2 | - | - |
| | Poitevine | 30 | - | - |
| | Provençale | 45 | - | - |
| | Provençale admixed with Alpine and Saanen | 11 | - | - |
| | Pyrénéenne | 30 | - | - |
| | Population from the German Alps admixed with other breeds[a] | 9 | - | - |
| | Appenzell | 5 | - | - |
| | Booted goat | 10 | - | - |
| | Capra grigia | 10 | - | - |
| | Grisons striped | 10 | - | - |
| | Nera verzasca | 10 | - | - |
| | Peacock goat | 10 | - | - |
| | Saanen | 30 | - | - |
| | Toggenburg | 10 | - | - |
| | Valais blackneck | 10 | - | - |
| | Local population from Italy | 10 | - | - |
| | Local population from Croatia | 22 | - | - |
| | Local population from Macedonia | 4 | - | - |
| | Angora | 17 | - | - |
| | Boer | 10 | - | - |
| **Total for wild type animals** | | **355** | **0** | **0** |
| Polycerate | Local population of Massif Central admixed with African dwarf goat[a] | - | 5 | - |
| | Local population of Normandy admixed with Nubian goat[a] | - | 5 | - |
| | Provençale | - | 31 | - |
| | Provençale admixed with Alpine and Saanen | - | 5 | - |
| | Rove admixed with Provençale | - | 4 | - |
| | Population from the German Alps admixed with other breeds[a] | - | 11 | - |
| | Local population from Italy | - | 13 | - |
| | Local population from Macedonia | - | 3 | - |
| **Total for polycerate animals** | | **0** | **77** | **0** |
| **Grand total** | | **355** | **77** | **0** |

**Supplementary Table 10. Details on caprine and ovine skull specimens considered in the morphometric analyses.** "ID" corresponds to the reference of the skull in its collection of origin. ENVA: Musée Fragonard, Ecole Nationale Vétérinaire de Maisons-Alfort, France; Halle: Domestic Animal Collections from the Central Natural Science Collections, Martin Luther University Halle-Wittenberg, Halle, Germany; INRAE: collection of INRAE UMR1313 GABI, Jouy-en-Josas, France; MNHN-ZM-AC: Collection d'anatomie comparée du Muséum National d'Histoire Naturelle, Paris, France; MZS: collection d'anatomie du Musée Zoologique de Strasbourg, France; ONIRIS : collection d'anatomie d'ONIRIS, Nantes, France. The horn formula provides information on the number and localisation of the horns on the left and right sides of the skulls, respectively, separated by a ";". Each "1" designate a horn while brackets indicate fusions between the bases of neighbouring horns. Year: Year of death or of entry in the collection. Juv.: juvenile individuals have between 6 and 12 months of age approximately. Ad.: Adult. Scan: method used for the digitization of the skull, with A: Artec Eva structured-light scanner, B: Breuckmann StereoScan structured light scanner, and P: photogrammetry (see Methods). ND: missing information.

| Phenotype, sex & species | Collection-ID | Scan | Horn formula | Breed or population | Year | Age |
|---|---|---|---|---|---|---|
| Polycerate male sheep | MNHN-ZM-AC-1845-266 | B | (111);11 | ND | 1845 | Ad. |
| | ENVA A-IV-N23 | B | 11;11 | Jacob | 1903 | Ad. |
| | ENVA A-IV-N25 | B | 11;11 | Jacob | 1903 | Ad. |
| | MNHN-ZM-AC-A-12126 | B | 11;11 | Local population from Tunis, Tunisia | 1884 | Ad. |
| | MNHN-ZM-AC-A-12127 | B | 11;11 | | 1884 | Ad. |
| | MNHN-ZM-AC-A-12128 | B | 11;11 | | 1884 | Ad. |
| | MNHN-ZM-AC-A-12130 | B | 11;11 | | 1884 | Ad. |
| | MNHN-ZM-AC-A-12131 | B | 11;11 | | 1884 | Ad. |
| | MNHN-ZM-AC-A-12132 | B | 11;11 | | 1884 | Ad. |
| | MNHN-ZM-AC-A-12133 | B | 11;11 | | 1884 | Ad. |
| | MNHN-ZM-AC-A-12195 | B | 11;111 | | 1902 | Ad. |
| | MNHN-ZM-AC-A-12196 | B | 11;11 | | 1902 | Ad. |
| | MNHN-ZM-AC-ae781 | B | 11;11 | ND | ND | Ad. |
| | MZS-Mam-03721 | B | 11;11 | Local population from Erzurum, Turkey | 1910 | Ad. |
| | MZS-Mam-01566 | B | 11;11 | Unknown | 1907 | Ad. |
| | MZS-Az15 | B | 11;11 | Jacob | 2001 | Ad. |
| | MZS-Ovi015 | B | 11;11 | Jacob | 2003 | Juv. |
| | INRAE-Pachot_M4C_Haie | B | 11;11 | Jacob | 2017 | Ad. |
| | Halle-Ofswi1 | A | 11;11 | Indian fat-tailed | 1882 | Ad. |
| | Halle-Ofswi3 | A | 11;(11) | Indian fat-tailed | 1885 | Ad. |
| | Halle-Ofswi8 | A | 11;(11) | Indian fat-tailed | 1888 | Ad. |
| | Halle-Ofswi9 | P | 11;11 | Indian fat-tailed | 1888 | Juv. |
| | Halle-Oprs1 | A | 11;11 | Persian | 1899 | Juv. |
| | Halle-Osom72 | A | 11;11 | Somali | ND | Ad. |
| | Halle-Opru1 | A | 11;11 | Local population from Peru | 1883 | Ad. |
| | Halle-Omsr13 | A | (111);11 | Local population from Masuria, Poland | 1894 | Ad. |
| | Halle-Opmm1 | A | 11;11 | Pommeranian sheep | 1909 | Ad. |
| | Halle-O20b | A | 11;11 | German sheep | 1888 | Ad. |
| | Halle-Omf183 | A | (11);(11) | Mufflon cross | 1889 | Ad. |
| | Halle-Our53 | P | (11);(11) | Urial x Rambouillet | 1891 | Juv. |
| Polycerate female sheep | INRAE-Pachot_F4C | B | 11;11 | Jacob | 2017 | Ad. |
| | Halle-Ojak2 | A | 11;11 | Jacob | 1995 | Ad. |

**Supplementary Table 10, continued**

| Phenotype, sex & species | Collection-ID | Scan | Horn formula | Breed or population | Year | Age |
|---|---|---|---|---|---|---|
| Wild type male sheep | MNHN-ZM-AC-1909-4 | B | 1;1 | Local population from Tunis, Tunisia | 1909 | Ad. |
| | MNHN-ZM-AC-1904-200 | B | 1;1 | ND | 1904 | Ad. |
| | MNHN-ZM-AC-1945-88 | B | 1;1 | Corsican mouflon | 1945 | Ad. |
| | MNHN-ZM-AC-2000-438 | B | 1;1 | Corsican mouflon | 1989 | Ad. |
| | MNHN-ZM-AC-A-12151 | B | 1;1 | Corsican mouflon | ND | Ad. |
| | MNHN-ZM-AC-A-12157 | B | 1;1 | Local population from Algeria | 1852 | Ad. |
| | MNHN-ZM-AC-A-12177 | B | 1;1 | Scottish blackface | 1891 | Ad. |
| | MNHN-ZM-AC-A-12187 | B | 1;1 | Local population from Gascony, France | 1902 | Ad. |
| | MNHN-ZM-AC-A-12188 | B | 1;1 | Local population from France | ND | Ad. |
| | MNHN-ZM-AC-A-12193 | B | 1;1 | Local population from France | 1884 | Ad. |
| | MNHN-ZM-AC-A-12194 | B | 1;1 | Local population from the Morvan Massif, France | 1884 | Ad. |
| | MNHN-ZM-AC-Astrakan2 | B | 1;1 | Astrakhan | ND | Ad. |
| | ONIRIS-ENVN_Ouessant | B | 1;1 | Ouessant | 1990 | Ad. |
| | Halle-Ofswi7 | A | 1;1 | Indian fat-tailed | 1886 | Ad. |
| | Halle-Okar22 | A | 1;1 | Karakul | 1923 | Ad. |
| | Halle-Omf116 | A | 1;1 | Mouflon cross | 1888 | Ad. |
| | Halle-Omf138 | A | 1;1 | Mouflon cross | 1887 | Ad. |
| | Halle-Omsr15 | A | 1;1 | Mauritian | 1895 | Ad. |
| | Halle-Ongr11 | A | 1;1 | Negretti | 1882 | Ad. |
| | Halle-Orbag288 | A | 1;1 | Rambouillet x Argali | 1906 | Ad. |
| | Halle-Oshl10 | A | 1;1 | Shetland | 1904 | Ad. |
| | Halle-Osom71 | A | 1;1 | Somali | 1924 | Ad. |
| | Halle-Okar153 | A | 1;1 | Karakul | 1930 | Ad. |
| | MNHN-ZM-AC-A-12173 | B | 1;1 | Astrakhan | 1884 | Ad. |
| Wild type female sheep | MNHN-ZM-AC-A-12179 | B | 1;1 | Local population from Iceland | 1891 | Ad. |
| | MNHN-ZM-AC-A-12224 | B | 1;1 | Local population from Mycenae, Greece | 1887 | Ad. |
| | INRAE-Pachot_F2C | B | 1;1 | Jacob | 2017 | Ad. |
| | MNHN-ZM-AC-A-12172 | B | 1;1 | Astrakhan | 1884 | Ad. |
| | Halle-O14 | A | 1;1 | Local population from Germany | 1899 | Ad. |

**Supplementary Table 10, continued**

| Phenotype, sex & species | Collection-ID | Scan | Horn formula | Breed or population | Year | Age |
|---|---|---|---|---|---|---|
| Polycerate male goats | MNHN-ZM-AC-1875-630 | B | 11;11 | ND | 1875 | Ad. |
| | MNHN-ZM-AC-1906-141 | B | 11;11 | ND | 1906 | Ad. |
| | MNHN-ZM-AC-A-12122 | B | 11;11 | ND | 1884 | Ad. |
| | INRAE-Bresson_M4C | B | 11;11 | Provençale X Alpine | 2017 | Ad. |
| | MZS-Mam-03727 | B | 11;11 | Unknown, animal raised in the Zoological Garden of Strasbourg | 1896 | Ad. |
| | Halle-Csom3 | P | 11;11 | Somali x German spotted goat | 1913 | Juv. |
| | Halle-CSom2 | A | 11;11 | Somali x German spotted goat | 1914 | Ad. |
| Polycerate female goats | Halle-C10 | A | 11;11 | German white goat | 2013 | Ad. |
| | Halle-C11 | A | 11;11 | German spotted goat | 2013 | Ad. |
| | Halle-CohneNr10 | A | 11;11 | German goat | ND | Juv. |
| | MNHN-ZM-AC-A12124 | B | 11;11 | ND | <1884 | Ad. |
| | Halle-Cbz40 | P | 11;11 | crossbreed#: Bezoar multicross | 1991 | Juv. |
| Wild type male goats | MNHN-ZM-AC-1884-2144 | B | 1;1 | ND | 1884 | Ad. |
| | MNHN-ZM-AC-1905-276 | B | 1;1 | ND | 1905 | Ad. |
| | INRAE-Bresson_M2C | B | 1;1 | Provençale X Saanen, France | 2017 | Ad. |
| | ONIRIS-ENVN_PIS | B | 1;1 | Saanen | 1990 | Ad. |
| Wild type female goats | Halle-C15 | A | 1;1 | House goat, Germany | ND | Ad. |
| | Halle-C16 | A | 1;1 | House goat, Germany | ND | Ad. |
| | Halle-C19 | A | 1;1 | House goat, Germany | ND | Ad. |

**Supplementary Table 11. Definition and designation on each side of the skull of the anatomical landmarks and sliding semi-landmarks used in the analyses.** Landmarks placement is illustrated in **Suppl. Fig. 11.**

| **Landmarks** | | **Definition** |
|---|---|---|
| **left** | **right** | |
| **s0** | **s5** | Most dorsal point of the *Foramen infraorbitale* |
| **s1** | **s6** | *Processus lacrimalis caudalis* (Process on Margo supraorbitalis of the lacrimal bone) |
| **s2** | **s7** | Most dorsal point of the lateral *Foramen supraorbitale* |
| **s3** | **s8** | Most caudal point of the fronto-zygomatic suture (*sutura frontozygomatica*) |
| **s4** | **s9** | Caudo-lateral point of the temporo-zygomatic suture (*sutura temporozygomatica*) |
| **s10** | **s11** | Most caudal point of the horizontal plate (*lamina horizontalis*) of the palatine bone |
| **s12** | **s13** | Most caudal point of the *Margo interalveolaris*, connection with the first premolar tooth |
| **s14** | **s64** | Neck of the cornual process (*collum processus cornualis*) |
| **s15 to s63** | **S65 to s113** | Points s15-s63 (resp. s65-s113) were placed round the suture line between the *processus cornualis* and the frontal bone, starting with point s14 (resp. s64) and turning in a dextral rotation (resp. sinister rotation) from the orbit and towards the back of the skull. Point s63 (resp. s113) overlays with point s14 (resp. s64) to close the loop.<br>When the animal had four separated horns, landmarks were positioned for the two horns located on the same side as follows: the frontal half circle of the upper horn was made between landmarks s14 (resp. s64) and s29 (resp. s79) ; the latter being ventral on the upper horn and closest to the lower horn. Then landmark s30 (resp. s80) was placed just opposite, on the lower horn (at the most dorsal level of it). Landmarks s31 (resp. s81) to s50 (resp. s100) were positioned on the lower horn, turning forward and downward, then coming back up. Landmark s50 (resp. s100) was set to landmark s30 (resp. s80) to close the loop. Then Landmark s51 (resp. s81) was positioned on landmark s29 (resp. s79) of the upper horn and the last half circle was completed on the caudal part of the upper horn with landmarks s51 (resp. s101) to s63 (resp. s113). |
| **s114** | **s115** | Most medial point of the fronto-nasal suture (*sutura frontonasalis*) |

**Supplementary Table 12. List of four-horned polycerate male sheep genotyped for the causative mutation and phenotyped for the distance between the lateral horns on the left side (dlhl) and for the distance between the upper horns (duh).**

| No | Breed or population | Heterozygous Polycerate | | Homozygous Polycerate | | Total |
|---|---|---|---|---|---|---|
| | | Dlhl ≤ duh | Dlhl > duh | Dlhl ≤ duh | Dlhl > duh | |
| 1 | Jacob | 11 | - | 1 | 6 | **18** |
| 2 | Manx Loaghtan | 2 | - | - | 4 | **6** |
| 3 | Ouessant admixed with Jacob | 2 | - | - | - | **2** |
| 4 | Ovino Quadricorna | 2 | - | - | 1 | **3** |
| | **Total** | **17** | **0** | **1** | **11** | **29** |

**Supplementary Table 13. Details on whole genome sequences considered in the present study. Ovine and caprine control individuals collected for the present analyses originate from all around the world.** a) 401 individuals from the original dataset were not considered : one Navajo-Churro and two Tibetan animals with unknown phenotypes were eliminated because of the occurrence of polycerate animals in these breeds, as well as 398 animals of unknown or composite breeds. b) These animals belong to the Chinese Sishui Fur sheep breed.

| Species | NCBI Bioproject or url | File format | Nb of cases | Nb of controls |
|---|---|---|---|---|
| Goat | www.goatgenome.org/vargoats_data_access.html | VCF | - | 1160 |
| | PRJEB39341 | FASTQ | 1 | - |
| **Total for goat** | | | **1** | **1160** |
| Sheep | PRJEB6025 | VCF | - | 180 |
| | PRJEB6495 | VCF | - | 14 |
| | PRJEB9911 | FASTQ | - | 3 |
| | PRJEB14098 | FASTQ | - | 7 |
| | PRJEB14418 | FASTQ | - | 24 |
| | PRJEB15642 | VCF | - | 10 |
| | PRJEB23437 | VCF | - | 99 |
| | PRJEB31241 | VCF | - | 535[a] |
| | PRJEB31930 | FASTQ | - | 6 |
| | PRJEB32110 | FASTQ | - | 26 |
| | PRJEB35553 | FASTQ | - | 2 |
| | PRJEB35682 | FASTQ | - | 20 |
| | PRJEB37460 | FASTQ | - | 3 |
| | PRJNA624020 | FASTQ | 10[b] | 250 |
| | PRJEB39341 | FASTQ | 1 | - |
| **Total for sheep** | | | **11** | **1179** |

**Supplementary Table 14. Information on transgenic mouse strains and details on PCR primers used for genotyping purpose.** Mut: amplicon encompassing Chr2 g.74,768,587_75,133,794del (mutant allele) ; Wt: segment encompassing the proximal breakpoint of Chr2 g.74,768,587_75,133,794del (wild type allele).

| Name of the strain | Name in original publication | Reference | Primer sequences |
|---|---|---|---|
| $Hoxd1^{Lac}$ | $Hoxd1^{tm1Ddu}$ | Zákány *et al.*, 2001 | GAGTTTCTCTTTGCTGTAATGAAGAGCT<br>TCACATTCTCCACGGGCAAGCC |
| $BAC^{HoxD}$ | $TgBAC^{HoxD}$ | Schep *et al.*, 2016 | ACAGCTGCCTCTGTGGCCTC<br>ATTACGCCAGCTGGCGAAAGGG |
| $BAC^{Mtx2}$ | - | This work | ACAGCTGCCTCTGTGGCCTC<br>ATTACGCCAGCTGGCGAAAGGG |
| $HoxD^{Del(151kb)lac}$ | Del(tpSB2-attP) | Andrey *et al.*, 2013 | ACTAGCCAGATCCAATGGACC<br>CTATTACGCCAGCTGGCGAAAGG |
| $HoxD^{Del(365kb)}$ | - | This work | GCCTGCACCTATGCAGTTTGAAAGG<br>CTCACAGAGTTCCTTAAACACTCCGAG (Mut)<br>CTGTTGAGTACATCCTATCATCAGGAGC<br>CTCAAAGTTGGGAGAAAGCAACAGTGC (Wt) |

**Supplementary Table 15. Details on PCR primers used for confirming the nucleotide sequence at the fusion points of variant g.115,652,290_116,155,699delins137kb.** See **Supplementary Fig. 3** for a scheme of the segments involved in the mutation and the corresponding fusion points identified with letters A to D.

| Fusion point | Primer sequences |
|---|---|
| A | GTTTAAAAGGTGGGGGAAGG/GTTTCAGGCATGCAACACAG |
| B,C,D | CATCCTGCAGCCTGTGAGTA/GGTGTCATCCCCTGTCTCAG |

**Supplementary Table 16. Details on primers used for quantitative RT-PCR analyses.**

| Species | Name of gene and primer pairs | Primer sequences | Amplicon size in bp |
|---|---|---|---|
| Sheep | OAR_HOXD1_exon2 | GTTGGCTATCTCGATGCGTC<br>GATCCGCACGAATTTCAGCA | 100 |
| Sheep | OAR_HOXD1_intron1 | GCTCCTTCCGGCAATTTTCT<br>CGGTTTGGGTCTTATGGGGA | 107 |
| Goat | CHI_HOXD1_exon2 | CCCTTCCCGTTCCCTTTTCT<br>GACGCATCGAGATAGCCAAC | 101 |
| Sheep | OAR_HPRT1 | GCCACCCATCTCCTTCATCA<br>TGCTGAGGATTTGGAGAAGGT | 94 |
| Goat | CHI_HPRT1 | TGGACTAATTATGGACAGGACCG<br>TATAGCCCCCCTTGAGCACA | 101 |
| Sheep | OAR_H2AFZ | TAAAGCGTATTACCCCTCGTCA<br>CACCACCAGCAATTGTAGCC | 90 |
| Goat | CHI_H2AFZ | GCGTATTACCCCTCGTCACTTG<br>CAGCAATTGTAGCCTTGATGAGA | 80 |
| Both | GAPDH | CACTACCATGGAGAAGGCTGG<br>GTGGTTCACGCCCATCACA | 106 |
| Both | YWHAZ | GGAGCCCGTAGGTCATCTTG<br>CTCGAGCCATCTGCTGTTTTT | 85 |
| Both | RPLP0 | TCTCCTTCGGGCTGGTCAT<br>AGGAAGCGGGAATGCAGAGT | 100 |

### Supplementary Note 1

**Absence of homozygous mutants at variant g.115,652,290_116,155,699delins137kb amongst 77 polycerate goats.**

Despite a continuous selection for polyceraty in most of the sampled herds, homozygous mutants were never observed amongst 77 polycerate animals for variant g.115,652,290_116,155,699delins137kb. Based on breeders' records, 14 of these goats were born from two polycerate parents while the other individuals were born from wild type x polycerate mating or from unknown sire with polycerate dam (i.e. when several mature males of different phenotypes are present in the same herd). Assuming that all the parents were heterozygous for variant g.115,652,290_116,155,699delins137kb, a Mendelian transmission would have generated 33% (i.e. 25%/(50% + 25%)) of homozygous mutant animals amongst the 14 polycerate goats born from polycerate parents. Considering a binomial law with parameters n=14 and p=0.33, the probability of not observing any homozygote is: $3.4 \times 10^{-3}$.

This strong presumption of homozygous lethality was supported by the analysis of a 365-kb deletion in mouse (g.74,768,587_75,133,794del on Chr2; murine genome assembly mm10), largely overlapping the orthologous 503-kb genomic segment absent in polycerate goats (g.115,652,290_116,155,699delins137kb on Chr2; **Fig. 1** of this note). Genotyping this variant at birth, in 42 animals derived from heterozygous parents did not identify any homozygous condition for the mutant allele, while at least 25% were expected (binomial $p=5.7 \times 10^{-6}$ with parameters n=42 and p=0.25; **Table 1** of this note).

*Mtx2* is the only protein-coding gene affected by the two deletions and morpholino knockdown of this gene in zebrafish is lethal at gastrulation (Wilkins et al. 2008). Altogether, these results indicate that the lack of *Mtx2* causes early embryonic death, which explains the absence of live homozygous mutants amongst the caprine and murine panels studied.

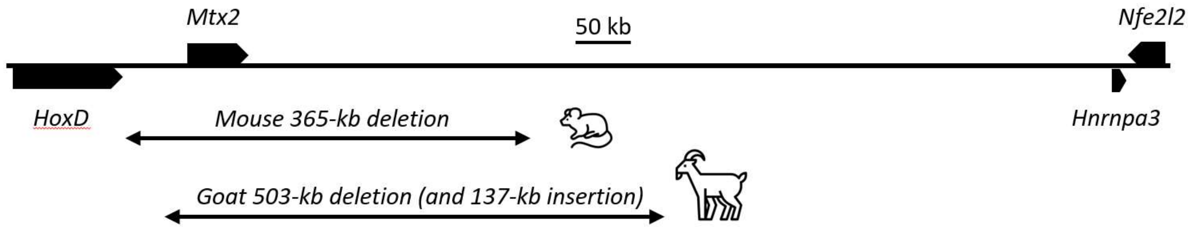


**Fig. 1 of Suppl. Note 1. Details on the 365-kb deletion encompassing *Mtx2* in mouse.** Murine and caprine orthologous segments of chromosome 2 with relative localizations of mouse g.74,768,587_75,133,794del and goat g.115,652,290_116,155,699delins137kb variants. Mouse and goat icons were made by "Monkik" from www.thenounproject.com.

| Number of animals per genotype | | | |
| --- | --- | --- | --- |
| Wt/Wt | Wt/Del | Del/Del | Total |
| 12 | 30 | 0 | **42** |

**Table 1 of Suppl. Note 1.** Results of the genotyping at birth of 42 mice born from mating between animals that were heterozygous for a 365-kb deletion encompassing *Mtx2*.
